# NAc-DBS selectively enhances memory updating without effect on retrieval

**DOI:** 10.1101/2025.01.16.633319

**Authors:** Andrés Pérez-Segura, Jorge Medina, Antonio Cerdán Cerdá, Cristina Sánchez-Ferri, Daniel Torres, Claudio R. Mirasso, Bryan Strange, Víctor M. Eguíluz, Lucas Lacasa, Santiago Canals

## Abstract

Deep brain stimulation (DBS) has emerged as a widely used therapeutic option when pharmacological treatments prove ineffective or refractory for psychiatric patients. The nucleus accumbens (NAc) represents a frequently targeted site in DBS interventions due to its demonstrated safety profile and therapeutic efficacy in obsessive-compulsive disorder, major depression, and anorexia nervosa. However, limited mechanistic understanding hampers its broader clinical applicability. This study sought to delineate the distinct behavioural dimensions modulated by NAc-DBS, its impact on distinct facets of memory, and to elucidate the underlying brain-network mechanism of action. We developed a novel spatial navigation task for rats and employed a high-dimensional behavioural analysis complemented by fMRI to dissect the cognitive, behavioural and neurobiological effects of NAc-DBS. Active NAc-DBS produced a selective enhancement of long-term memory encoding without affecting memory recall or working memory. We found no effect of NAc-DBS on motor, appetitive or stress-related behaviours. Sustained neuronal activation in the NAc, septum, entorhinal and insular cortex demonstrated no desensitization to chronic NAc-DBS, which triggered a functional reorganization among dopaminergic-related structures. These findings suggest that NAc-DBS induces a functional reorganization in the mesocorticolimbic system, potentially mimicking a dopaminergic novelty signal to enhance memory updating. This provides a mechanistic basis for the therapeutic use of NAc-DBS, particularly in improving cognitive flexibility in psychiatric disorders.

## INTRODUCTION

Neuromodulation has emerged as an effective therapeutic approach for psychiatric disorders when pharmacological treatments prove ineffective or refractory for patients^1,2^. Among these techniques, deep brain stimulation (DBS) stands out for its precise spatial and temporal control of neural activity. The nucleus accumbens (NAc), a structure in the basal forebrain with a central role in reward, pleasure and reinforcement learning, is a widely used DBS target in medication-resistant cases of obsessive-compulsive disorder, major depression or anorexia nervosa^3-5^. More generally, the profound capacity of DBS to alter neural activity posits it as a potentially transformative tool for enhancing cognitive capacities.

In a recent study in patients with obsessive-compulsive disorder receiving DBS in the NAc-medial septum, we found enhanced mnemonic function during stimulation^6^. Indications of improved cognitive and memory scores were also described in other DBS studies targeting the NAc in humans^7,8^. In animal models, several studies have implicated the NAc in the formation of hippocampal-dependent memories^9-10^, and our previous research demonstrated that blocking NAc activity disrupts the functional reorganization of memory-related brain regions driven by long-term synaptic plasticity^11^. We hypothesized that the symptomatic improvements observed in psychiatric conditions may result from enhanced memory updating facilitated by NAc neuromodulation, which, in turn, promotes increased cognitive flexibility.

Despite its widespread use, the mechanism of action of DBS remains incompletely understood. Many processes and mechanisms may come into play, from the psychological through to the neurobiological. Does DBS affect attention, memory encoding, memory retrieval or other cognitive processes? And, mechanistically, how does the composition of the stimulated tissue or the electrical characteristics of the electrode and the stimulation protocols influence brain activation^12,13^?. Identifying which brain regions will be modulated by direct or indirect action of stimulation, and in which direction, is a key issue of interest. Factors such as local *vs*. remote activity modulation by either orthodromic and/or antidromic recruitment, whether *en passant* axons are activated, or what synaptic plasticity will come into play in prolonged stimulation protocols, all contribute to this complexity. To better understand DBS mechanisms, the use of animal models with translational value is essential. Animal models enable the recording and quantification of stimulation effects at the neuronal circuit level and allow for associations with cognitive-level observations.

The present study aims to elucidate the effects of DBS in the NAc on memory formation and memory recall. A new behavioural task was used, modelled in part on previous research^14^, that is capable of dissecting the potential impact of DBS on these two aspects of memory. We also integrated high-dimensional behavioural analysis, enriching cognitive readouts in a rat model, along with functional magnetic resonance imaging (fMRI) to assess DBS-driven changes in brain activity. Memory studies in animal models often rely on learning curves and basic metrics such as event counting, exploration times, or times to task completion. These approaches, as in loss- and gain-of-function experiments^15-18^, evaluate the impact on memory encoding, consolidation, or retrieval separately, often overlooking potential confounding factors such as attentional or motor effects. In this study, by dissecting the behavioural effects of DBS using a multi-dimensional analysis and uncovering the neural networks modulated in both the short and long term, we approach a more comprehensive understanding of how NAc-DBS influences memory processes, potentially guiding future neurotherapeutic interventions.

## MATERIALS AND METHODS

### Experimental design

A total of 37 animals were used in four different experiments. Three behavioural experiments utilised 25 male Long Evans rats purchased from Janvier labs (France), weighing around 250 g at the beginning of the experiments. They were paired and housed in a controlled environment with a room temperature of 21±2 °C, following an inverted light cycle where lights were on from 20:00 to 8:00. This setup allowed behavioural procedures to be conducted during the rats’ dark cycle. Throughout the experiments, the rats had *ad libitum* access to food and water.

The first experiment was aimed to validate the newly developed behavioural task (n = 6). The second experiment was devoted to perform DBS of the NAc (n = 4). From an initial group of 18 rats we selected for each group those exhibiting greater exploratory behaviour to proceed with the experiments. This selection was made over the first two days of habituation by excluding animals that spent most of their time close to the walls of the arena, as subsequent memory analysis would rely on characterizing the exploratory behaviour of the animals. The integrity of the implants for DBS during the whole duration of the extensive behavioural training and tests conditioned the permanence of the animals in the study. These two experiments comprised a total of 95 behavioural experimental tests, including DBS sessions of three linked sessions (S1-3, see *The navigation task* section) in a within-subject manner of active (current on) and sham (current off) stimulation, which were statistically compared and increased the statistical power of the comparison. The two conditions were counterbalanced for each animal.

The third behavioural experiment involved the evaluation of possible anxiolytic or appetitive effects of the NAc-DBS treatment (n = 7). In conditioned place preference (CPP) experiments all rats underwent DBS in the less preferred chamber for each rat (see Supplementary Information). Elevated plus maze (EPM) was performed by randomly interleaving DBS-ON and -OFF periods during the exposure of the animals to the maze.

The fourth experiment was a fMRI investigation of brain networks modulated by DBS. The number of repetitions of each stimulation paradigm used is specified below. We utilised 12 male Sprague-Dawley rats, purchased and housed under similar conditions.

All experiments were conducted in compliance with both Spanish and European legislation.

### Deep brain stimulation

DBS was administered bilaterally continuously throughout the entire execution of the behavioural tasks on selected days. The stimulation protocol, which was designed to match those applied to humans in the clinic, involved biphasic pulses beginning with the negative phase, delivered at a frequency of 130 Hz. Each pulse had a total duration of 120 μs and an amplitude of 250 μA (Fig. 3A). The stimulation intensity was also adjusted to match the voltages commonly used in clinical procedures^1-3,6,19^. Trials with and without stimulation (sham trials) were randomly counterbalanced. In sham trials, animals were connected to the stimulation setup with current set to 0.

For the fMRI experiments unilateral stimulation was used in order to discern contralateral activations as well, and three distinct stimulation paradigms were employed. To assess the acute effect of stimulation, a block-designed protocol involving 10 stimulation trains were utilised. The total duration of each train (ON-period) was 8 seconds, followed by an OFF-period of 22 seconds. The same stimulation paradigm, delivered without off-periods in 30-minute trains (4 trains), was employed to investigate the effects of continuous DBS stimulation. The effects of the DBS cessation, designed to reveal a possible inhibitory effect of the stimulation that is released upon cessation of stimulation, were assessed with the application of blocks of 10-minutes trains performing fMRI acquisitions during the end of those events (12 events acquired, Fig. 4B).

### The navigation task

The task takes place within a circular arena measuring 180 cm in diameter, featuring four entering/exit boxes situated at the North, East, South, and West positions. The doors of these entering boxes are operated by pneumatic pistons. The apparatus is housed in a temperature-controlled room set at around 21 °C, provided with visual cues on the walls outside the maze. A recording camera is positioned above the maze (Flea3, Teledyne FLIR, USA), along with a dimmable light source. The recorded image is analysed and processed in real-time to control the maze based on the tracking of the animal’s behaviour. All components of the apparatus are integrated into an Arduino UNO board (Arduino LCC) and orchestrated using the Bonsai visual reactive programming software^20^, synchronised with online animal tracking.

In the task, upon entering the arena through one of the boxes, the animal navigates under high illumination conditions (approximately 400 lux) to locate a user-defined virtual (invisible for the rat) platform. Once the rat locates the virtual platform, the light switches off and the entering door reopens, allowing the animal to return to the box. In this dry version of the Morris water maze, the natural aversion of rats to intense light serves as both an aversive stimulus and a cue to signal when the “platform” has been reached, motivating the animals to actively seek for it (Fig. 1A). In regular experiments, the virtual platform has a diameter of 12 cm and can manifest in any of the 16 predetermined locations (see Fig.1B).

**Fig. 1.**
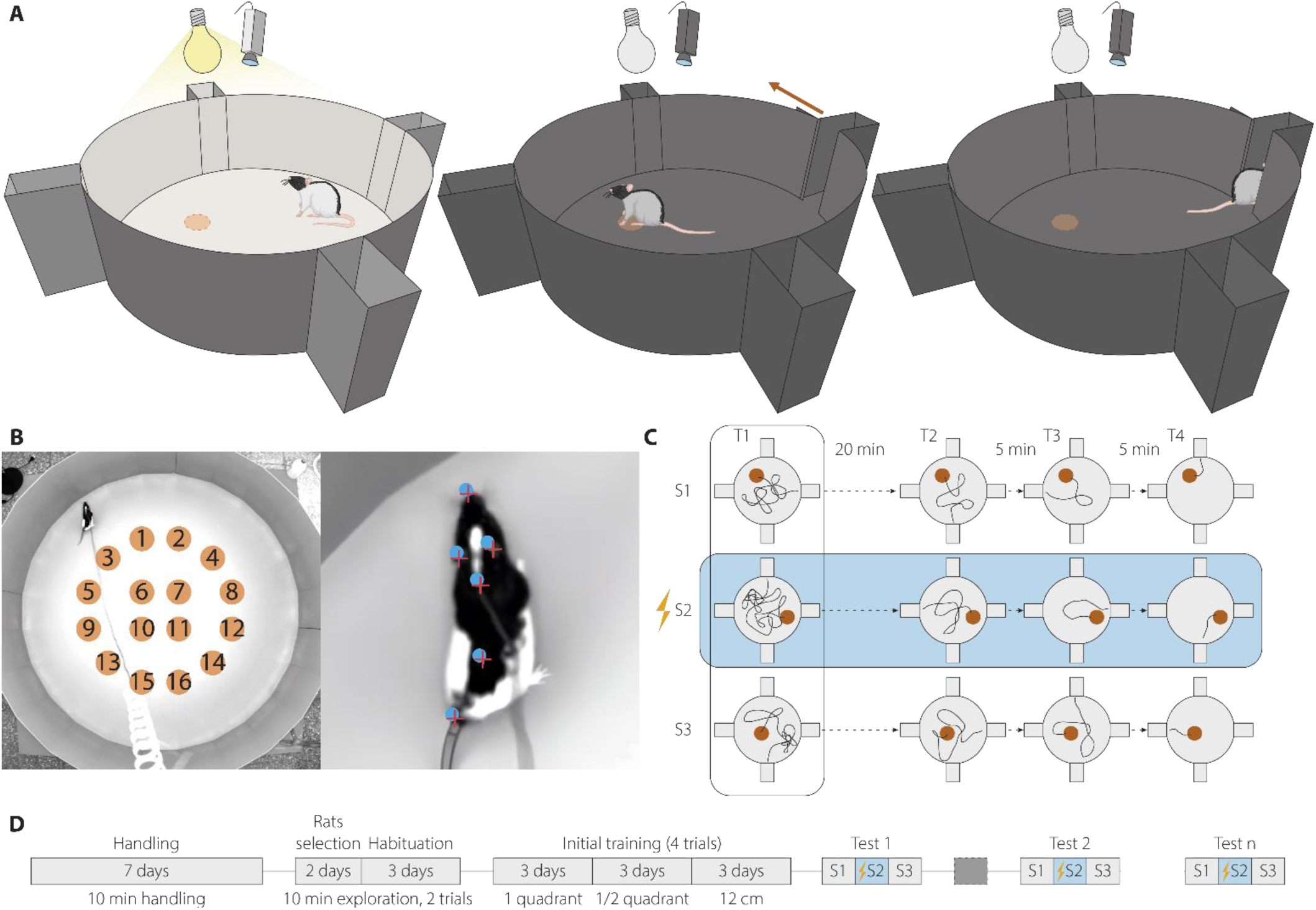
The spatial navigation task. **(A)** Representation of the circular arena (180 cm diameter) with the four entering boxes. The animal navigates in a brightly illuminated arena (400 lux) to find a computer-defined but invisible virtual platform (dotted orange circle). When the animal reaches the “platform” the illumination dims and the exit door open allowing the animal to return to the entering box. **(B)** Image of the real maze from the top with the 16 possible virtual platform locations. In the right panel, a rat with the body parts used for tracking labelled (nose, left and right ears, shoulders, back and base of the tail). **(C)** Schematic of the three consecutive sessions (S1→S3), one per day, of a typical memory experiment, and the four trials performed each session for measuring and proper encoding the new “platform” location. The location of the “platform” is selected between the 16 preselected locations pseudo randomly to avoid repeating locations across days. Experimental manipulation (NAc-DBS) on the second day (S2, highlighted in blue) allows investigating memory retrieval, encoding and updating (see STAR Methods). **(D)** Training schedule. After 7 days of handling and 5 of habituation, the animals completed 9 days of training to learn the task. Sequences of experimental sessions (S1→S3) were then intercalated with sessions in which the virtual platform does not change its location between days (dark grey box). Animals may repeat as many sequences of sessions as required.

One week prior to commencing training, animals undergo 7 days of handling, spending 10 minutes daily within the experimentation room under comfortable light conditions. Subsequently, they undergo a 5-day habituation protocol. During the first two days, the animals freely explore the arena without illumination by being placed directly inside the arena. Over the next three days, they enter the arena from randomly selected entering boxes, intensively illuminated, while the arena remains dark. Once the door opens, the animals explore the arena for 10 minutes, repeating this task twice daily with a 20-minute delay. The initial two days of habituation help identify animals that spend more time in the central area, ensuring a group with highly exploratory individuals (Fig. 1D).

In the initial training phase, the animals learn the contingencies of the task (a specific location in the large arena switches off the light and opens an escape pathway to conclude the task). This phase comprises three stages over three consecutive days, starting with large virtual platforms and progressively reducing their size. The first stage starts with a one-quadrant size in the centre of the arena, followed by half a quadrant, and concludes with the final size of the “platform”. Animals are considered trained at this point. Figure 1D provides a schematic representation of the complete schedule.

A specific feature of the design allows dissociation of memory formation and memory recall and adapts the delay-matched-to-place protocol^14,17^ previously developed for the water maze. The animals are extensively trained in this task for 15 sessions which are, importantly, structured in successive groups of 3 sessions called S1-S3 (Fig 1C). An experimental session in one session consists of 4 trials (T1-4) of up to 10 min in which the virtual platform remains in the same position (Fig. 1C). The location of the virtual platform changes the next day, and it is pseudo randomly moved between the 16 locations across successive sessions. Thus, on T1, the animals may remember the location of the virtual platform of the previous session one day earlier; however, across trials T1 to T4, they will encode a new location and so form a new memory. This new location may be remembered on T1 of the next day. Accordingly, during the first 90 sec of T1, the virtual platform is inactive to allow precise monitoring of the searching behaviour of the animals. By having successive groups of 3 sessions together, DBS could be applied on S2 to examine its impact on recall of memory formed on S1, its impact on short-term memory during S2, and its impact on the formation of long-term memory measured on T1 of S3 (Fig. 1C).

Memory function is estimated by extracting different measures from the tracking of the animals (see *Behavioural data preprocessing* section) related to the search for the previous day’s region of interest (ROI, enlarged version of the virtual platform with a diameter of 24 cm, long term memory, LTM) and how it adapts to the new virtual platform location (short term memory, STM). Non-experimental test experiments with no manipulation and maintaining the “platform” location between days, occur between triads of experiments to keep the animals engaged in the task.

### Behavioural data processing

The Bonsai visual reactive programming software^20^ was employed for online tracking to operate the apparatus based on the animal’s location in the arena. The received camera image underwent processing with an HSV threshold to generate a binary image, facilitating the detection of when the animal reached specified locations.

For data analysis, precise tracking of the animals (nose, left and right ears, shoulders, back, and the base of the tail, Fig.1B) was extracted using the freely accessible software DeepLabCut^21,22^ (see Supplementary Information). Due to visual differences between groups (wired head implants in the DBS experiment), two distinct models were trained for data analysis—one for the non-stimulated animals and another for the DBS and control conditions. The diameter of the ROI for detecting exploration in the subsequent analysis was increased 2 times in order to prevent slight tracking imprecisions and to compensate for the distance between the shoulders (the main body part used in the analysis) and the silhouette of the animal (used to operate the apparatus).

In the navigation task, the evidence of STM is derived from the exploration of the ROI in the 4 consecutive trials of a given day. For each trial, we meticulously analysed the navigation patterns of the animals towards the target. A comprehensive set of 20 metrics designed to delineate various aspects of navigation was extracted, which fell into 5 main categories: directionality towards target, wall navigation strategies, ballisticness, spatial navigation, and speed (see Table 1). The variation of these metrics between trials 1 (T1) and 4 (T4) concerning the current ROI location was used to evaluate STM. Subsequently, LTM is quantified by statistically comparing the variation of previously described behavioural metrics in the T1 of a given day concerning the current ROI and the previous day’s ROI. The median statistic is employed to penalise outliers and one-tailed confidence intervals with confidence level 95 % were then used to determine significance (see Supplementary Information).

**Table 1.**
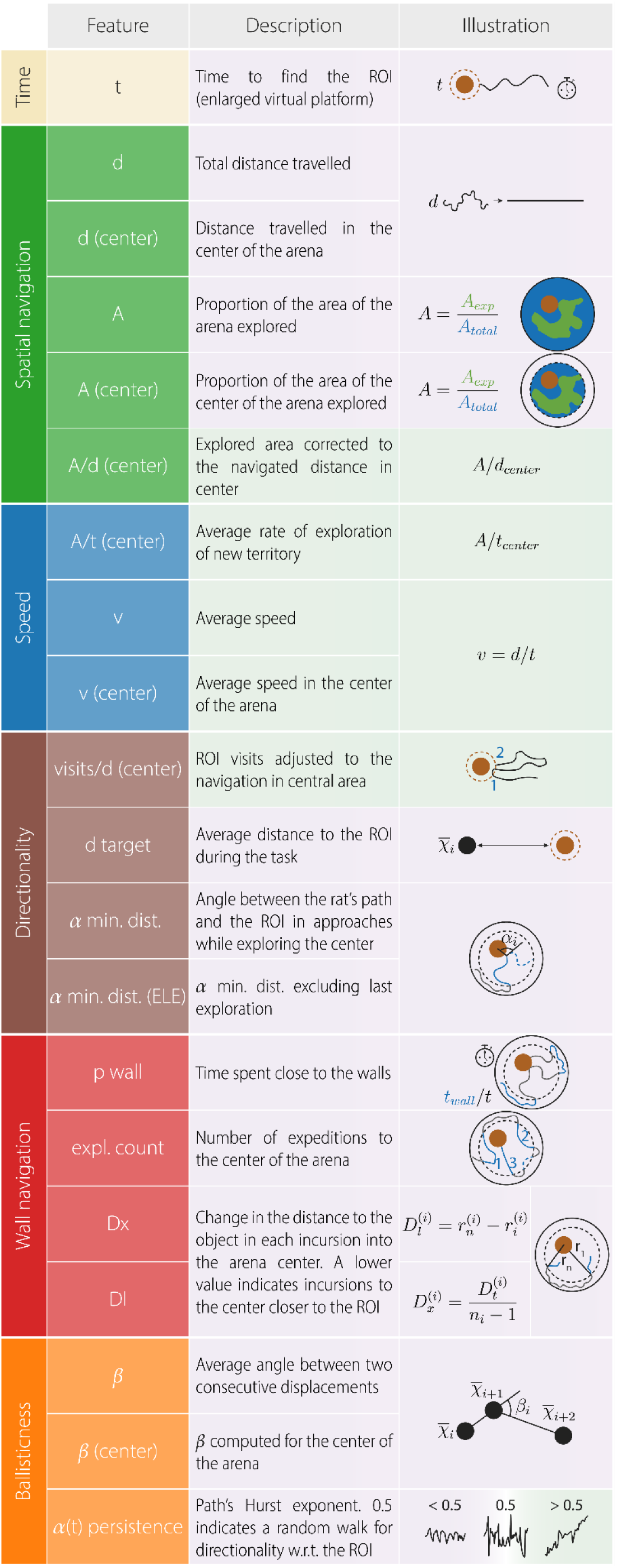
Behavioural features. Representation of the metrics used to quantify STM and LTM. Each metric assesses different aspects of the rats’ navigation and is related to categories such as time (yellow), spatial navigation efficiency (green), speed (blue), directionality (brown), wall navigation strategies (red) and ballisticness (orange). Features on the purple or green background are expected, within this behavioural task, to decrease or increase, respectively, when navigation is guided by a more precise memory representation of the virtual platform location.

|  | Feature | Description | Illustration |
| --- | --- | --- | --- |
| Time | $t$ | Time to find the ROI (enlarged virtual platform) | |
| Spatial navigation | $d$ | Total distance travelled | |
| | $d$ (center) | Distance travelled in the center of the arena | |
| | $A$ | Proportion of the area of the arena explored | $A = \frac{A_{exp}}{A_{total}}$ |
| | $A$ (center) | Proportion of the area of the center of the arena explored | $A = \frac{A_{exp}}{A_{total}}$ |
| | $A/d$ (center) | Explored area corrected to the navigated distance in center | $A/d_{center}$ |
| Speed | $A/t$ (center) | Average rate of exploration of new territory | $A/t_{center}$ |
| | $v$ | Average speed | $v = d/t$ |
| | $v$ (center) | Average speed in the center of the arena | |
| Directionality | visits/$d$ (center) | ROI visits adjusted to the navigation in central area | |
| | $d$ target | Average distance to the ROI during the task | $\bar{x}_i$ |
| | $\alpha$ min. dist. | Angle between the rat's path and the ROI in approaches while exploring the center | |
| | $\alpha$ min. dist. (ELE) | $\alpha$ min. dist. excluding last exploration | |
| Wall navigation | $p$ wall | Time spent close to the walls | |
|  | expl. count | Number of expeditions to the center of the arena |  |
| | $D_x$ | Change in the distance to the object in each incursion into the arena center. A lower value indicates incursions to the center closer to the ROI | $D_l^{(i)} = r_n^{(i)} - r_i^{(i)}$ |
| | $D_l$ | | $D_x^{(i)} = \frac{D_l^{(i)}}{n_i - 1}$ |
| Ballisticness | $\beta$ | Average angle between two consecutive displacements | $\bar{x}_{i+1}$ $\beta_i$ $\bar{x}_{i+2}$ |
| | $\beta$ (center) | $\beta$ computed for the center of the arena | |
| | $\alpha(t)$ persistence | Path's Hurst exponent. 0.5 indicates a random walk for directionality w.r.t. the ROI | |

The original set of 20 metrics was primarily designed to evaluate the presence of STM and LTM (first-order effects). The selection of second-order metrics, used to assess differences in memory performance between DBS and control groups, was pre-determined based on their ability to measure first-order effects (see Supplementary Information). Additionally, a decision model was applied to detect LTM. If a rat exhibits LTM, it will tend to explore the ROI from the previous day first during T1, rather than the new, unknown ROI. This defines a null model for the absence of LTM: a binary classifier predicting that the rat will first locate the target closest to its entry point into the arena. If this null model performs worse than the proportion of rats finding the previous day’s ROI first, the null model is rejected, and LTM is confirmed (see Supplementary Information).

All calculations for the offline analysis were performed in Python and other standard statistical procedures were computed in SPSS (IBM Corp. Released 2022. IBM SPSS Statistics for Windows, Version 29.0) and GraphPad Prism (GraphPad Software, Version 8).

## RESULTS

To investigate the effect of DBS in the NAc on memory formation, we developed a behavioural navigation task inspired in the delayed-matching to place protocol of the Morris water maze^17^. This new task, and its detailed analysis, takes into account navigation trajectories, their directionality, persistence, biases and tendencies towards walls and central zones, as well as present and past target locations (see Materials and Methods). Collectively, these help to differentiate effects of DBS on memory encoding and retrieval in a single intervention.

### Spatial memory task

In a large and brightly illuminated circular arena (diameter 180 cm), rats are trained to find a virtual platform with its successful localization serving to dim the intense lights and open an escape door (Fig. 1A). The circular maze has 4 entrance/exit doors, and the entire operation is automated and guided in real time by video tracking. The virtual platform is a circular region of 12 cm diameter randomly placed in one of 16 predefined locations in the maze (Fig. 1B).

A specific feature of the design allows dissociation of memory formation and memory recall. The animals are extensively trained in this task for 15 sessions which are, importantly, structured in successive groups of 3 sessions called S1-S3 (Fig. 1C). After a period of habituation and training (Fig. 1D, Materials and Methods), an experimental session in one day consists of 4 trials (T1-4) of up to 10 min in which the virtual platform remains in the same position (Fig. 1C). The location of the virtual platform changes the next day and is moved between the 16 locations across successive sessions. Thus, on T1, the animals may remember the location of the virtual platform of the previous session one day earlier; however, across trials T1 to T4, they will encode a new location and so form a new memory. This new location may be remembered on T1 of the next day. Accordingly, during the first 90 sec of T1, the virtual platform is inactive to allow precise monitoring of the searching behaviour of the animals. By having successive groups of 3 sessions together, DBS could be applied on S2 to examine its impact on recall of memory formed on S1, its impact on STM during S2, and its impact on the formation of LTM measured on T1 of S3 (Fig. 1C). For offline behaviour quantification purposes, we defined a ROI with a diameter of 24 cm, centred on the virtual platform used in each session (see Materials and Methods).

Once the initial training is completed (Fig. 1D, 2A), the duration for the animal to reach the ROI diminishes progressively across the four daily trials (repeated measures ANOVA Geisser-Greenhouse corrected, F(trial)_1.96, 147.7_ = 6, p = 0.003. Fig. 2B). This result underscores the animals’ understanding of task contingencies and allows quantification of STM within the session. The navigation of the animal on the following day (S2) towards the ROI location encountered on the previous day (S1) was, as noted, used to investigate LTM (Fig. 1C). The reason the searching time is long on T1 across multiple sessions is because the animals are remembering the location of the virtual platform of the previous day.

**Fig. 2.**
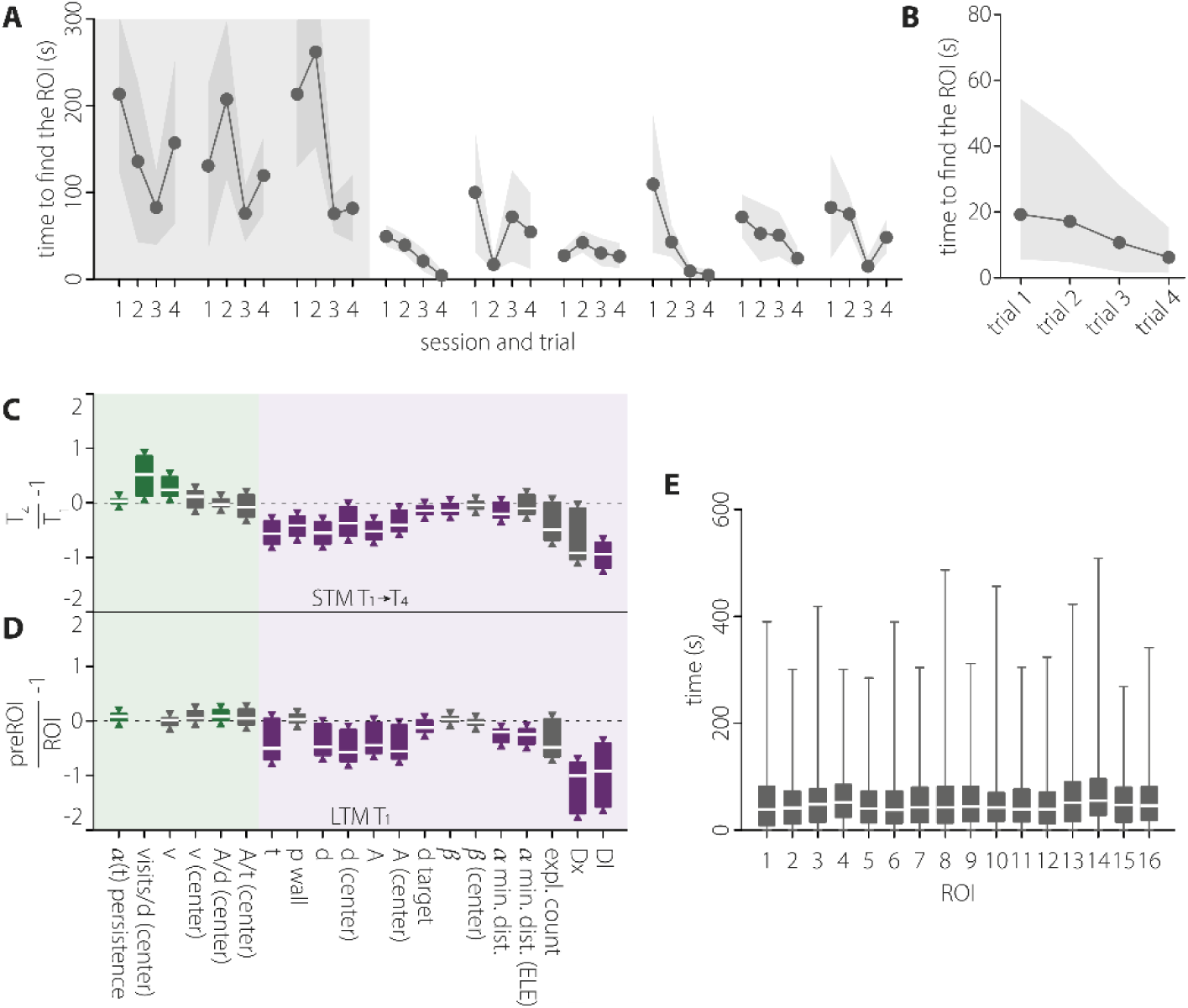
Memory evaluation in the navigation task. **(A)** Quantification of the time needed to find the ROI (an enlarged version of the virtual platform used to operate the maze, see STAR Methods) of the rats across the last stage of the initial training (grey background) and the first six sessions, representing a typical performance. n = 6, mean ± SEM. **(B)** Quantification of the time needed to find the ROI in each trial performed after the completion of the initial training. 6 rats * 15 sessions, median ± interquartile range. **(C)** Quantitative evidence for STM. Equally tailed, 95 % CI for the median ratio change between T1 and T4. Metrics on light green or light purple background are expected to adopt more positive or negative values, respectively, when the memory representation of the virtual platform (ROI) location improves. Green and purple-coloured boxplots indicate a statistically significant differences at the 95 % confidence level (one tailed interval). **(D)** Same as in C for the quantitative evidence for LTM, measuring with respect to the ROI and the previous day’s ROI (preROI) in T1 across sessions. **(E)** Average time to find all possible ROI locations in all experiments. Median and min-to-max whiskers represented, n = 2910.

For each trial, we meticulously analysed the navigation patterns of the animals towards the target. A comprehensive set of 20 metrics delineating various aspects of navigation was extracted, which fell into 5 main categories: directionality towards target, wall navigation strategies, ballisticness, spatial navigation, and speed (see Table 1 and Materials and Methods for details). Comparing animal trajectories between T1 and T4 of all sessions, unveiled statistically significant changes in 13 of these metrics after correcting for multiple comparisons, controlling for a false discovery rate (FDR) of 0.05. The joint evidence supports the acquisition of the new location into STM (Fig. 2C, Table S1). Notably, along the four trials, rats exhibited a more directed and less erratic navigation towards the target as indexed by the parameters *ß* and *α(t) persistence* (ballisticness), searched repeatedly in the correct area (*visits/d(center)*) and performed paths more directed to the virtual platform (*α min. dist.*), navigated closer to the target (*d target*) with less distance navigated, both in the central zone of the arena (*d(center)*) and overall (*d*), and needed to cover less area of the arena to find the target (*A* and *A(center)*, Fig. 2C). Probably as a consequence of the more efficient and directed exploration, animals found the target earlier (*t*, Fig. 2C). Moreover, the rats developed a navigation strategy characterised by circumnavigation around the walls to approach to the target ROI from a shorter distance to the wall, a behaviour quantified by the *Dl* metric (Fig. 2C). This strategy was accompanied by an increase in movement speed (*v*), increasing the overall proportion of time spent near the walls (*p wall*, Fig. 2C).

We then sought to determine whether any inherent biases could influence rats’ navigation towards specific regions of the arena, such as more centric *vs.* peripheral ROIs. We quantified the time required for rats to locate each potential ROI in every trial following the initial training phase. Our analysis revealed no significant differences in the time needed to find the various ROIs (Kruskal-Wallis test, χ^2^ = 23.59, p = 0.07, df = 15, Fig. 2E).

We next assessed the existence of LTM by analysing the navigation trajectories during the first trial (T1) on the following day (S2) towards the ROI that was active the previous day (S1). We compared the rat’s tendency to navigate towards the previous day’s ROI (S1) *versus* navigation directed to the current (still unknown) ROI (S2) or any other potential ROI location in the maze. It was observed that rats were more likely to reach first the S1’s ROI on T1 of S2 than the current ROI (time ratio to reach S1 or S2 ROIs in T1 of S2, P(*t*_S1ROI_ < *t*_S2ROI_) = 0.63, Table 2, see Materials and Methods). This result could not be explained by random navigation, as indicated by the rejection of the null hypothesis stating that arrival is due to chance based on ROI proximity to the entering door (H_0_: *t*_closest_ < *t*_furthest_ rejected, Table 2, see Materials and Methods), not being the closest ROIs the more likely to be reached first.

**Table 2.**
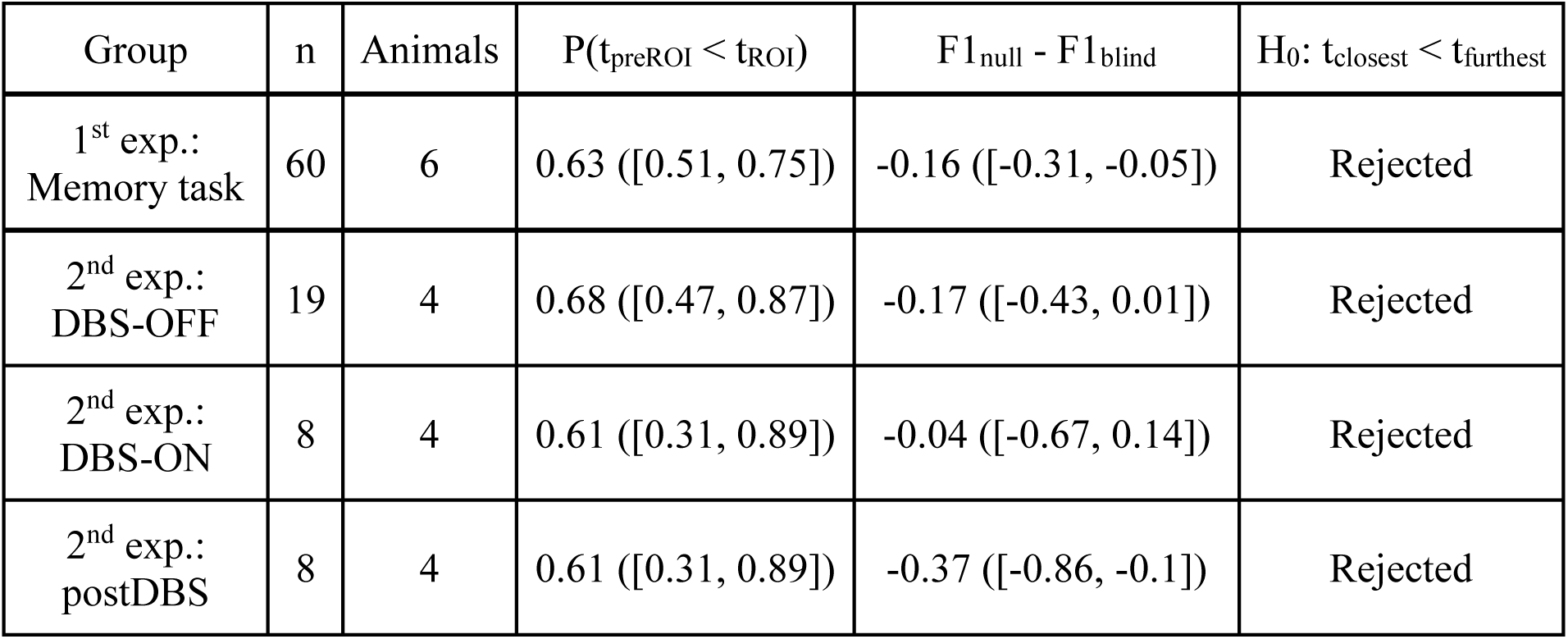
LTM evaluation. Experiments (group), observations (n), number of animals and metrics for arrival times used to evaluate LTM. P(*t*_preROI_ < *t*_ROI_) is the ratio of sessions finding before the ROI in the previous session (preROI) than the current ROI. This behaviour was not solely due to proximity to the starting position (F1_null_ - F1_blind_, H_0_: *t*_closest_ < *t*_furthest_; see STAR Methods). The postDBS group refers to animals receiving NAc-DBS on S2, with LTM assesed on T1 of S3.

| Group | n | Animals | $P(t_{\text{preROI}} < t_{\text{ROI}})$ | $F1_{\text{null}} - F1_{\text{blind}}$ | $H_0: t_{\text{closest}} < t_{\text{furthest}}$ |
| --- | --- | --- | --- | --- | --- |
| 1<sup>st</sup> exp.:<br>Memory task | 60 | 6 | 0.63 ([0.51, 0.75]) | -0.16 ([-0.31, -0.05]) | Rejected |
| 2<sup>nd</sup> exp.:<br>DBS-OFF | 19 | 4 | 0.68 ([0.47, 0.87]) | -0.17 ([-0.43, 0.01]) | Rejected |
| 2<sup>nd</sup> exp.:<br>DBS-ON | 8 | 4 | 0.61 ([0.31, 0.89]) | -0.04 ([-0.67, 0.14]) | Rejected |
| 2<sup>nd</sup> exp.:<br>postDBS | 8 | 4 | 0.61 ([0.31, 0.89]) | -0.37 ([-0.86, -0.1]) | Rejected |

Importantly, rats also demonstrated in S2 significantly more directed navigation towards the S1 ROI (Fig. 2D, Table S2), taking less time to reach it (*t*, Fig. 2D), constraining their exploration trajectories to the area surrounding that ROI (*α min. Dist*., *α min. Dist. (ELE)*, *d target*, Fig. 2D) and tracing more ballistic trajectories (*α(t)* persistence, Fig. 2D) with less spatial redundancy (*A/d(center)*, Fig. 2D). Additionally, their exploration of the arena enroute to the previous session’s ROI was markedly more efficient, minimising the distance travelled, and the area covered before reaching the virtual platform (*d*, *d(center)*, *A*, *A(center)*, Fig. 2D). As for STM, rats exhibited in T1 the same circumnavigation strategy around the walls to approach the previous session’s target ROI from the shortest distance to the wall (*Dx*, *Dl*, Fig. 2D).

Information stored in LTM is progressively updated by the new information encountered in the arena over trials T1-T4, as searching first the previous ROI in T1 (significance level α = 0.05) was substituted by searching first at the current ROI in T4 (significance level α = 0.075).

### NAc-DBS experiments

The previous experiments confirmed the existence of STM and LTM formation, both detectable within our experimental setup. Next, we examined the impact of NAc-DBS on memory formation over 4 successive series of three linked sessions (S1-3) in a within-subject manner, which increases the statistical power of the comparison. Control stimulation (DBS-OFF) sessions were included, with DBS-ON and -OFF sessions counterbalanced for each animal. Stimulation was applied bilaterally and continuously during the whole duration of each of the 4 trials of session S2 (see Fig. 3A and Materials and Methods for stimulation details).

**Fig. 3.**
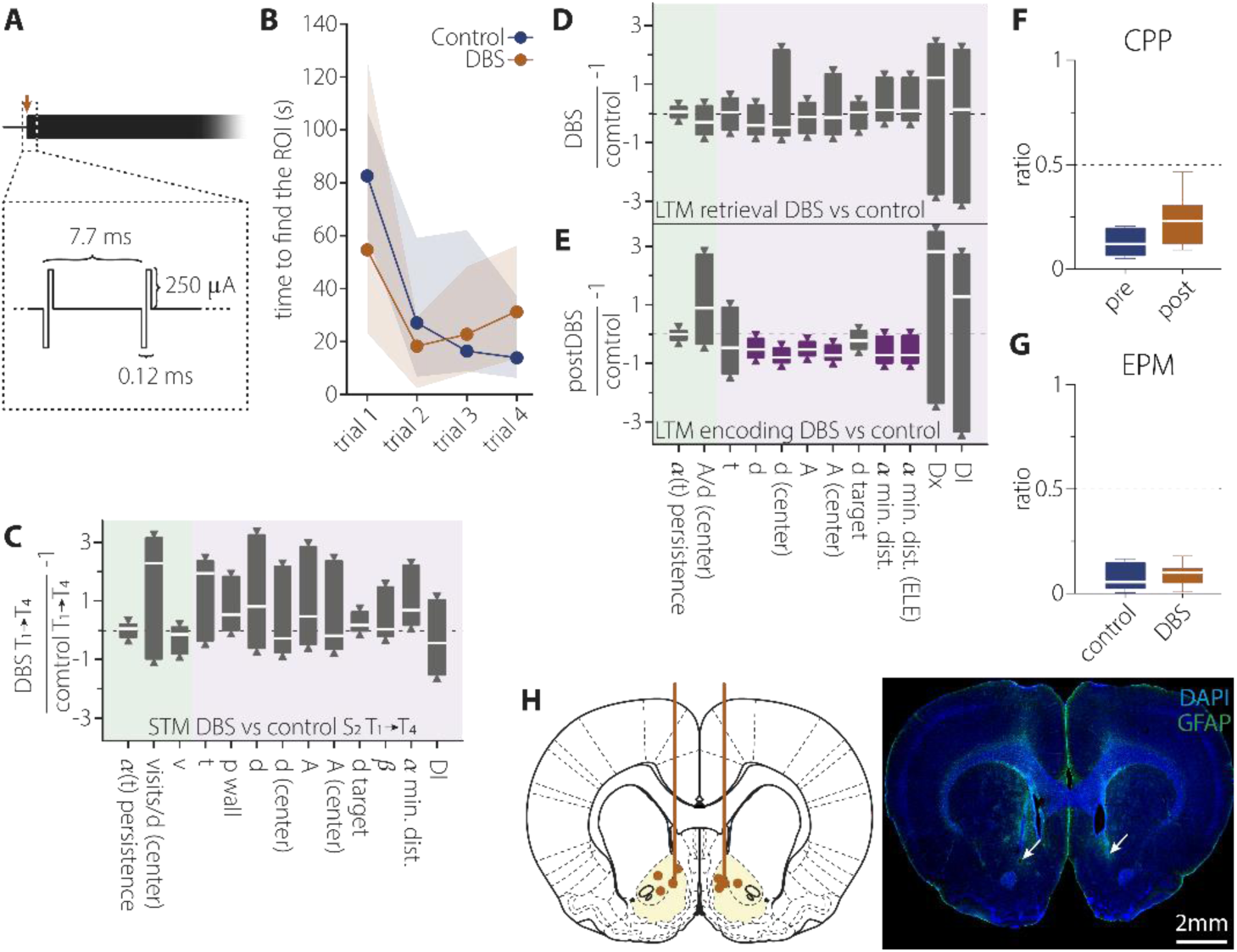
Effect of DBS in the NAc on memory performance. **(A)** Schematic representation of the DBS protocol, consisting of 250 μA biphasic pulses at 130 Hz delivered bilaterally and continuously during the 10 min of each trial of session S2 in selected experiments. **(B)** Quantification of the time to find the ROI in each trial in S2, in DBS-OFF (Control) and DBS-ON (DBS) conditions. 10 sessions for DBS-OFF and 8 sessions for DBS-ON, median ± interquartile range. **(C)** Effect of DBS on STM performance. Equally tailed 95 % CI for the median ratio of metrics comparing DBS-OFF (Control) *vs.* DBS-ON (DBS) in T4. Metrics on green or purple background are expected to adopt more positive or negative values, respectively, when the memory representation of the platform location improves. Green and purple-coloured boxplots indicate a statistically significant differences at the 95 % confidence level (one tailed interval). **(D)** Same as C for the effect of DBS on LTM retrieval, measuring at T1 of S2 the searching of the previous day’s ROI in sessions with DBS-OFF (Control) *vs.* DBS-ON (DBS). **(E)** Same as D for the effect of DBS on LTM encoding, measuring at T1 of S3 the searching of the ROI that was encoded the previous day during DBS-ON (postDBS) *vs.* DBS-OFF (Control). **(F)** Effect of DBS on CPP. Preference is measured as the ratio of the time spent in each chamber with respect the stimulated one. DBS is applied in the less preferred chamber for each subject. Median and min-to-max whiskers represented, n = 7. **(G)** Effect of DBS on EPM. Exploration ratio between the open divided by closed arms, during DBS-OFF (Control) vs. DBS-ON (DBS) periods. Median and min-to-max whiskers represented, n = 7. **(H)** Schematic representation of the implanted electrodes with the tip of the electrodes marked by orange dots. Lower panel shows a representative example of a histological preparation. White arrows indicate the tip of the electrodes.

Changes in STM and LTM induced by NAc-DBS are considered second-order effects with respect to the existence of STM and LTM (which would be the first-order effects). From a Bayesian perspective on metric design, we argue that the prior probability of a metric detecting a change in STM and LTM (the second-order effect) depends on its ability to detect the first-order effect. Consequently, the metrics designed to capture changes in STM and LTM in the NAc-DBS experiment are limited to the subset of the original 20 metrics that showed significant first-order effects. Metrics unable to detect such second-order effects cannot inform the study’s conclusions.

#### NAc-DBS effects on short-term memory

During STM formation, an analysis of arrival times to the ROI under both DBS-ON and DBS-OFF conditions, revealed a significant decrease between T1 and T4 trials in both groups, with neither stimulation effect or interaction between trials and conditions reaching significance (Mixed-effects model analysis F(trial)_1.5, 23.58_ = 6.28, p(trial) = 0.01, F(condition)_1, 16_ = 0.35, p(condition) = 0.7, F(interaction)_3, 47_ = 1, p(interaction) = 0.4, Fig. 3B). Consistent with the analysis on arrival times, high-dimensional behavioural analysis, performed on the subset of metrics validated for their ability to detect STM (Fig. 2C, Table S1), showed no significant differences that passed multiple comparison corrections between DBS-ON and DBS-OFF conditions. Overall, these findings indicate that NAc-DBS does not significantly facilitate or interfere with spatial STM formation.

#### NAc-DBS effects on long-term memory retrieval

We next assessed the effect of NAc-DBS on the retrieval of previously encoded LTM. We found that animals during DBS-ON sessions retain the ability to find the previous day ROI (S1) earlier than the current ROI (S2) at T1 of S2 (P(*t*_S1ROI_ < *t*_S2ROI_) = 0.611, H_0_: *t*_closest_ < *t*_furthest_ rejected, Table 2). The behavioural analysis, performed on the subset of metrics validated for their ability to detect LTM (Fig. 2D, Table S2), showed no difference between DBS-ON and OFF conditions (Fig. 3D). These results demonstrate no positive nor negative interference of NAc-DBS on the retrieval of previously acquired memories.

#### NAc-DBS effects on long-term memory encoding

We then investigated the effect of NAc-DBS on LTM encoding by assessing target navigation on T1 of S3. This is, we assessed, in the absence of stimulation (S3), the retrieval of memories encoded the previous day (S2) in the presence (DBS-ON) or absence (DBS-OFF) of NAc stimulation. First, the probability to first reach the target confirmed the ability of the animals to find the ROI location encoded in S2 before than the current ROI location of S3 (P(*t*_S2ROI_ < *t*_S3ROI_) = 0.611, H_0_: *t*_closest_ < *t*_furthest_ rejected, Table 2). Importantly, animals that were stimulated in S2 (DBS-ON) showed more efficient navigation towards the S2 ROI on S3 than non-stimulated ones (Fig. 3E, Table S3), suggesting improved memory formation by NAc-DBS. Specifically, stimulated animals covered less distance and explored a smaller area to find the target (*d, d (center), A, A(center)*, Fig. 3E). Their movements were also more ballistic towards the target (*α min. Dist., α min. Dist. (ELE)*, Fig. 3E). These results remained significant after multiple comparison correction using the Benjamini-Hochberg method, controlling for a FDR of 0.1 (Table S3). Notably, with six significant results, the expected number of false positives is 0.6 i.e., between zero and one test. Thus, the conclusions drawn from the collective evidence remain robust following correction.

Overall, these findings suggest that DBS in the NAc may exert a favourable influence on the encoding phase of memory formation, without significantly disrupting either the retrieval of memories during the stimulation period or affecting other measured behavioural parameters to a notable extent.

#### Memory enhancement is unlikely to be linked to appetitive, anxiolytic, or motor effects

Given the extensive dopaminergic afferents in the NAc^23^, we investigated potential appetitive or anxiolytic effects of DBS that could hinder the interpretation of results.

For the appetitive effect, we used a CPP task by applying the NAc-DBS while the animal was in one of the 3 chambers of the apparatus (see Supplementary Information). No statistically significant differences were found between DBS-ON and OFF conditions (two-tailed paired t-test, t = 2.04, df = 5, p = 0.1, Fig. 3F). Similarly, anxiety evaluation was conducted using the EPM (see Supplementary Information) and revealed no significant effect of DBS. The proportion of time animals spent exploring the open arms of the maze did not significantly differ between DBS-ON and -OFF conditions (two-tailed paired t-test, t = 0.35, df = 6, p = 0.7, Fig. 3G).

Potential stimulation-induced motor alterations are unlikely, as demonstrated by the absence of significant differences in the animals’ navigation speed (*v*, *v(center)*) when exploring the arena in the first trial of S2 (LTM retrieval test) while the stimulation is ON *vs.* OFF. The tortuosity of trajectories (*α(t) persistence*) also did not change (Fig. 3D). We argue that a direct effect on motor function should be evident in all DBS-ON trials, which, as we indicated above, was not the case.

At the end of the behavioural experiments, the position of the DBS electrodes in the NAc was verified (Fig. 3H). Furthermore, there was no excessive tissue reaction to the implanted electrode, as assessed by astroglial staining.

### Changes in brain activity induced by NAc-DBS

Using MRI-compatible stimulation electrodes and recording fMRI signals during the application of the DBS protocol, we investigated changes in brain activity induced by the stimulation (Fig. 4). In this case, unilateral stimulation was used in order to discern contralateral activations as well. Acute stimulation of the NAc in blocks of 8 seconds (Fig. 1B, see Materials and Methods) evoked robust blood oxygenation level dependent (BOLD) signal responses and enabled us to map brain regions afferent and efferent to the stimulated area, including *en passant* axons. Major evoked responses by acute stimulation were found in the NAc and other striatal regions, the prefrontal, orbitofrontal, insular and entorhinal cortices, the septum and the ventral hippocampus (Fig. 4C and D).

**Fig. 4.**
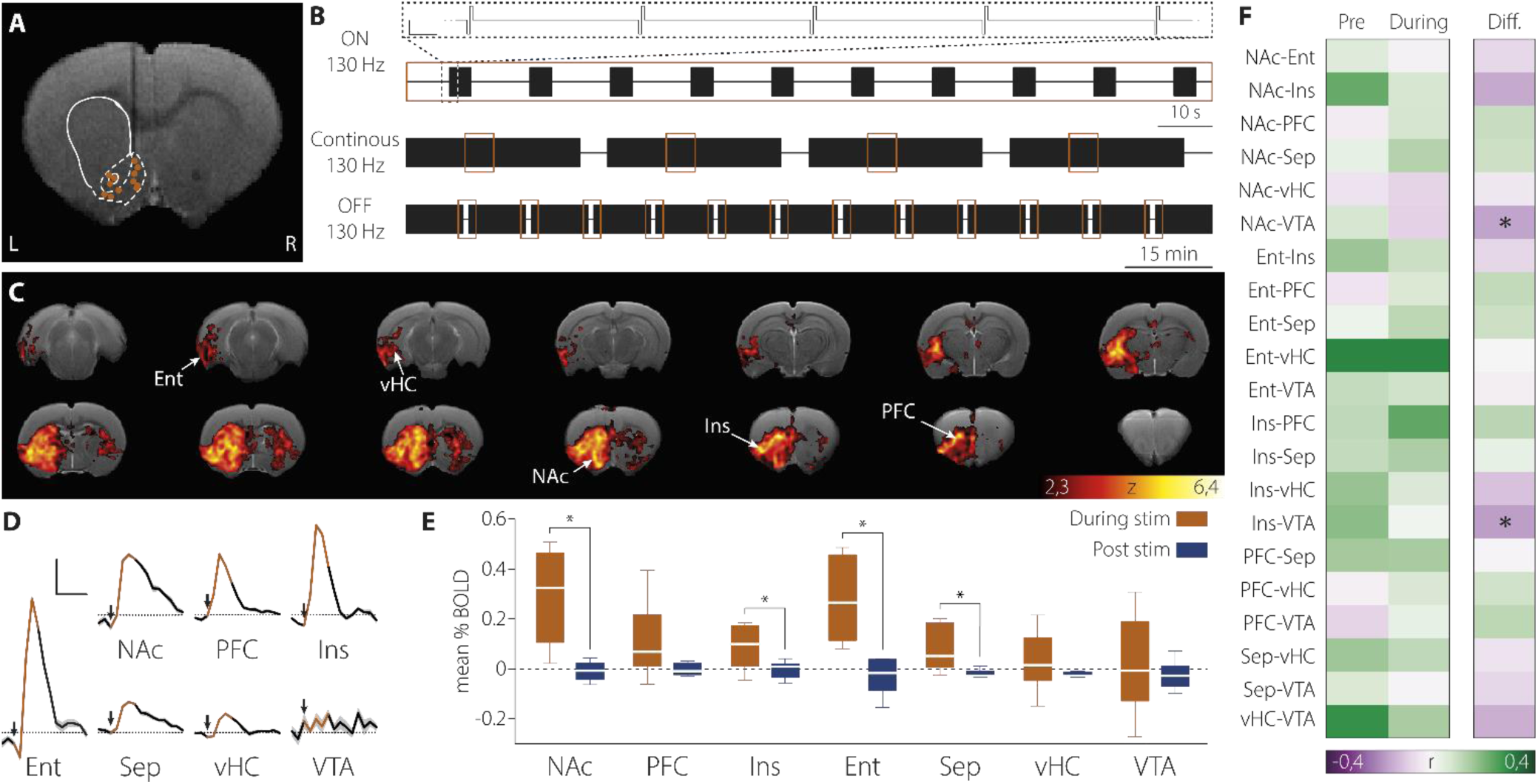
Acute and chronic effects of NAc-DBS on brain-wide activity. **(A)** High resolution anatomical image (T2-weighted) showing the location of the stimulation electrode in each animal (while lines representing the striatal area, with the core and shell parts of the NAc delimited with dotted lines). The vertical thin dark shadow is the artefact produced by the carbon-fibre DBS electrode. **(B)** Schematic representation of the stimulation protocols used. Top: acute DBS ON paradigm (8 s ON, 22 s OFF). Scale 1ms, 300 μA. Middle: continuous DBS paradigm. fMRI data (10 min) is recorded during continuous DBS stimulation. Bottom: acute DBS OFF paradigm. After a period of sustained DBS (10 min) the stimulus is switched off and the fMRI response is recorded. Orange squares represent acquisition windows. **(C)** Thresholded cluster-corrected functional maps (z > 2.33, cluster p = 0.05) of the animals stimulated with the acute DBS ON paradigm overlaid on a T2-weighted anatomical image. n= 12. **(D)** BOLD signal time courses evoked by the acute DBS ON in selected brain regions, represented as % change *vs*. baseline. Black arrows mark the start of the stimulation and orange tracing the duration of the stimulus. Scale 10 s, 0.5 %, n= 12. **(E)** BOLD signal changes in the acute DBS OFF paradigm. Orange bars: mean BOLD signal in the last 22 s of the 10 min continuous stimulation. Bue bars: mean BOLD signal in the first 22 s after stimulation cessation. Median and min-to-max whiskers represented, n = 6, *p < 0.05. **(F)** Pearson correlation between the mean BOLD signals in selected brain regions during continuous DBS *vs*. resting-state pre-stimulation (Pre) conditions. Right panels show the difference between both conditions n=4, *p(Benjamini-Hochberg corrected) < 0.05.) NAc: nucleus accumbens, PFC: prefrontal cortex, Ins: insular cortex, Ent: entorhinal cortex, Sep: septum, vHC: ventral hippocampus, VTA: ventral tegmental area.

To investigate the effect of DBS during a chronic application, as performed in the behavioural experiments above, we substituted acute by continuous stimulation and recorded fMRI signals in 10 min blocks. We then computed functional connectivity as the correlations between the average BOLD signals recorded in the regions identified in the previous experiment (Fig. 4D). We compared functional connectivity before and during DBS application (Fig. 4F). Significant changes in correlation between the measured brain regions were induced by the continuous stimulation (one-way MANOVA, F_21, 6_ = 4.17, p = 0.04, Wilks’ lambda = 0.064, Fig. 4F). Pairwise comparisons for the 21 pairs of brain regions revealed statistically significant differences between DBS-ON and OFF conditions between the ventral tegmental area (VTA) and the NAc (F_1, 26_ = 12.42, p(Benjamini-Hochberg corrected) = 0.02, FDR correction for the 21 comparison tests), and between the insular cortex and the VTA (F_1, 26_ = 20.85, p(Benjamini-Hochberg corrected) = 0.002, FDR correction for the 21 comparison tests), decreasing in both cases with NAc-DBS.

The direct effect of continuous electrical stimulation at high frequency on neuronal activity is not well understood. Arguments in favour of a neuronal inactivation in response to a prolonged stimulation can be found in the literature^12,13,24,25^. To investigate this hypothesis, we acquired fMRI signals during the last 22 seconds of a 10-min stimulation period and for the next 22 seconds after the end of stimulation (Fig. 4B). If neuronal response to DBS retains a certain level of activity in the continuous protocol, the BOLD signal is expected to decrease after stimulus termination. Conversely, if neuronal response is completely habituated to stimulation, no change in BOLD signal level is expected after the stimulus is switched off. Alternatively, if continued stimulation actively inhibits the target area, rebound activity would be expected due to disinhibition upon cessation of stimulation. We found that the mean BOLD signal significantly decreased upon DBS cessation in the NAc (two-tailed paired t-test, t = 3.66, df = 5, p = 0.01), entorhinal cortex (two-tailed paired t-test, t = 3.32, df = 5, p = 0.02), insular cortex (two-tailed paired t-test, t = 3.54, df = 5, p = 0.02) and septum (two-tailed paired t-test, t = 2.93, df = 5, p = 0.03), with no measurable rebound activity in any structure (Fig. 4E). These findings reveal that, under the current experimental conditions and throughout the 10-minute duration of the behavioural task, continuous DBS induced a sustained activity increase across several of the targeted structures, including the NAc.

## DISCUSSION

We provide evidence supporting a positive effect of DBS in the NAc on long-term memory formation. Specifically, we demonstrated that DBS can facilitate the encoding of new memories without affecting the retrieval of existing memories or interfering with ongoing short-term memory processes. This efficacy, combined with the absence of significant appetitive or anxiolytic side effects, underscores the considerable potential of NAc-DBS for clinical application. Our research also highlights the importance of employing high-dimensional behavioural analysis, which provided a more nuanced understanding of DBS facilitating mechanistic interpretations. Importantly, most of the NAc-DBS-driven effects on memory performance reported here would have otherwise gone unnoticed using standard "time-to-target" behavioural measurements. Lastly, by combining NAc-DBS and simultaneous whole-brain measurements with fMRI, we provide evidence for sustained *vs.* putatively phasic effects of chronic stimulation on distal brain areas.

### Learning in the newly developed navigation task

The procedure developed allowed effective memory encoding of each daily location, with performance typically characterised by a monotonic decrease in latency to find the ROI across trials in each session (Fig. 2B). The asymptotic performance in trials 2-4 reflects the effectiveness of the memory update. High-dimensional analysis showed that better memory formation was indicated by more effective navigation of the maze, covering a shorter distance and exploring a smaller area of the maze before finding the target, with more frequent and faster forays to the central zone being made from the wall position nearest to the target ROI, with directed and ballistic trajectories.

In each session, the animals first recalled the memory of the ROI’s previous location and subsequently updated their memory by encoding its current location within the session. The same set of high-dimensional features indicated STM updating with the new location of the ROI within each session (between T1 and T4) and demonstrated LTM between sessions (at T1 of S2 or S3, compare Figs 2 A and C). Overall, the behavioural task and analysis provide a robust framework for evaluating the effects of interventions on distinct facets of memory formation. This is achieved within a continuous task repetition paradigm, which helps mitigate the impact of small group sizes that are often challenging to increase due to the complexity of some animal preparations. By maximizing the data obtained from each subject, the framework also minimizes the number of animals required for a study.

### Dissecting NAc-DBS effects using the navigation task

We have shown that NAc-DBS can enhance LTM in rats. The NAc is implicated in the reward system and is integrated within the memory network^10^. Inactivation of the NAc in rats inhibits the functional reorganisation within the memory network associated with long-term synaptic potentiation in the hippocampus^11^, which otherwise enhances the functional coupling between memory related structures like the hippocampus and prefrontal cortex^26,27^. In humans, an improvement in episodic memory formation has been recently reported in a cohort of obsessive-compulsive disorder patients, and one patient with anorexia nervosa, receiving 130 Hz electrical stimulation in the NAc^6^. Our findings support this observation and provide neurobiological insight into the phenomenon. Additionally, given that human studies are conducted in psychiatric patients, our controlled animal study rules out any potential effect of DBS on improving symptoms of the disease that could confound, or at least contribute to the observed effects of NAc-DBS on memory in these patients.

NAc-DBS increases LTM performance, an effect it produces by facilitating the codification of new information rather than facilitating the recall of previously stored information, supporting a critical on-line role of this structure for memory enhancement^6^. This differentiation is crucial for interpreting the effects of DBS and planning its potential therapeutic use. Additionally, the absence of significant changes in the high-dimensional behavioural space analysed during LTM retrieval in DBS-ON vs. -OFF conditions dismisses the notion of unspecific enhancement of other cognitive or executive functions by NAc-DBS.

In a broader context, we propose that the facilitation of memory encoding by NAc-DBS may promote a more flexible memory system, enabling more efficient cycles of encoding, recall, updating, and consolidation of updated information. This enhanced flexibility could contribute to overall cognitive function and may underlie the therapeutic benefits of NAc-DBS in patients with obsessive-compulsive disorder and anorexia nervosa, conditions characterised by pronounced cognitive rigidity^3^. Furthermore, cognitive flexibilization driven by a more efficient memory updating process may explain why the effectiveness of NAc-DBS treatment in psychiatric patients is not immediate but instead improves progressively over the course of treatment^2^.

Finally, despite the dense reciprocal connectivity with the VTA and the importance of the dopaminergic system in appetitive behaviour and anxiety^10,23,28-30^, we observed only a minor trend, not reaching statistical significance, indicating DBS effects on appetitive behaviour, with no observable effect on anxiety. This lack of significant findings avoids complications in interpreting behavioural changes attributed to memory effects.

### Brain-wide interactions of NAc-DBS

Acute (8 sec long) stimulation in the NAc at 130 Hz engaged a network of memory-related brain regions extending from the NAc to the septum, insular, prefrontal and entorhinal cortices and the ventral hippocampus, in agreement with previous fMRI studies^31^. However, stimulation is delivered continuously in most DBS protocols including ours. In those conditions, brain activations in response to stimulation habituated in some degree, but retained supra-baseline levels in the NAc, septum, insular cortex and entorhinal cortex. Sustained activation of this network may explain why a reversible memory effect of NAc-DBS is observed in patients even after long periods of DBS^6^. Furthermore, continuous DBS decreased the functional connectivity between the VTA and the NAc, and between the VTA and the insular cortex.

Sustained activity in the NAc by induced DBS may involve continuous dopamine release from VTA axons, which could act as a saliency signal enhancing memory encoding^10^. Additionally, the activation by DBS of the efferent GABAergic connection from the NAc to the VTA^32,33^, composed of the axons of D1 medium spiny neurons targeting GABAergic interneurons in the VTA, may lead to a disinhibited state of the VTA and a further increase in dopaminergic release in the hippocampus and other memory-related structures^10,33^. This interpretation is supported by the transition from a positive to a negative correlation between the BOLD signals in the VTA and NAc (Fig. 4F). This disinhibited state would enhance the responsiveness of the VTA to inputs coming from different brain regions. A larger variety of inputs driving VTA activity may also account for the change in BOLD signal correlations between the VTA and the NAc and insular cortex (Fig. 4F). Our results support a long-lasting reorganisation of the functional balance in the mesocorticolimbic system, resulting in sustained enhanced activity in regions critical for memory formation.

When conducting fMRI experiments, the use of anesthesia necessitates caution in directly interpreting brain connectivity results to explain cognitive processes in awake animals. Although this limitation is widely acknowledged, it is also important to highlight some advantages of using urethane as an anesthetic agent. Urethane has been shown to induce brain states more akin to physiological sleep than to coma-like states^34^ and, unlike other commonly used anesthetics such as isoflurane, minimally impacts blood pressure and blood flow^35^. This characteristic is particularly relevant to our study, as fluctuations in these parameters could significantly influence the acquired BOLD signals. Additionally, urethane’s capacity to maintain a stable depth of anesthesia over extended periods^36^ proved especially beneficial for our experimental paradigm, which involved analyzing the evolution of whole-brain interactions under the cumulative effects of DBS treatment over time. Future studies implementing an awake animal preparation for DBS-fMRI will be instrumental in assessing the extent to which these findings translate to the unanesthetized state.

Overall, the combination of an animal model enabling the dissection of various stages of memory formation with a high-dimensional behavioural analysis has led us to conclude that NAc-DBS selectively facilitates the updating or encoding of new information into memory without exerting positive or negative effects on recall and without significantly impacting other cognitive or executive functions. DBS-fMRI experiments further demonstrated that chronic stimulation induces sustained activation of the mesocorticolimbic system, prominently involving the dopaminergic system, which we suggest emulates a dopaminergic novelty signal that enhances memory encoding. We further propose that the therapeutic benefit of NAc-DBS in various psychiatric conditions may result from enhancing cognitive flexibility by promoting an updated memory base.

## ACKNOWLEDGEMENTS AND FUNDING

We are grateful to Professor Richard G.M. Morris for his advice on the design of the behavioural test, as well as for his insightful comments on the manuscript. We also thank Victor J. Rodriguez, Analia Rico and Clara Serrano for their excellent technical support. We acknolwled support from the Spanish Ministerio de Ciencia e Innovación, Agencia Estatal de Investigación PID2021-128158NB-C21 (SC), PRE2019-087450 (APS), DYNDEEP EUR2021-122007 (LL), MISLAND PID2020-114324GB-C22 (LL), PID2021-128158NB-C22 (CRM) funded by MICIU/AEI/10.13039/ 501100011033. We also acknowledge the Program for Centres of Excellence in R&D Severo Ochoa CEX2021-001165-S (SC, APS) and Maria de Maeztu project CEX2021-001164-M (LL, CRM, JM), Maria de Maeztu Excellence Unit MDM-2017-0711 (JM) funded by the MICIU/AEI/10.13039/501100011033.

## DECLARATION OF INTERESTS

Authors declare that they have no competing interests.

## AUTHOR CONTRIBUTIONS

Conceptualization: SC, APS, ACC

Formal analysis: APS, JM

Funding acquisition: SC, LL, CRM

Investigation: APS, ACC, CSF, DT

Methodology: SC, BS, APS, CRM, ACC, JM, VME, LL

Project administration: SC

Software: APS, JM

Supervision: SC, LL

Validation: APS, JM

Visualization: APS

Writing – original draft: SC, APS

Writing – review & editing: SC, APS, LL, JM, BS

## SUPPLEMENTARY INFORMATION

### Supplementary methods

#### Tracking analysis

DeepLabCut software^21,22^ was used to extract the precise position of the selected body parts of the animals across each frame of the videos. Each condition (wired and non-wired animals) involved preparing two sets of 16 videos for training. For each set, 40 frames of each video were utilised to label desired body parts (nose, left and right ears, shoulders, back, and the base of the tail, Fig.1B). Subsequently, 95% of the labelled frames were used for training the network.

A ResNet-101-based neural network^37,38^ was employed with default parameters for 1.3 x 10^6^ training iterations and three consecutive training sessions. The results were validated through a single shuffle, yielding a test error of 3.95 pixels and a train error of 3.5 pixels for the control condition. In the DBS condition, the test error was 5.17 pixels, and the train error was 3.9 pixels. The image size used was 1552 x 1552 pixels. Subsequently, these networks were applied to analyse the remaining recorded videos for each respective group.

#### Long-term memory evaluation

For LTM detection and regarding the probability of a rat in finding the target ROI or any other ROI in the arena, we defined a null model (lack of LTM): a binary classifier predicting that a rat first finds the target (either ROI or previous day’s ROI) whose location is closer to the rat’s entering point to the arena:

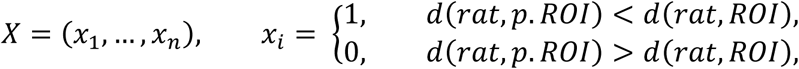

The null model is then evaluated against the actual outcome (finding first the previous day’s ROI or the ROI at each T1) across rats and days:

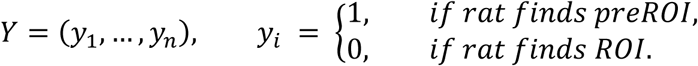

Lack of LTM suggests that this null model should outperform a blind classifier that simply predicts the majority class (without using any data). To compare both classifiers, the F_1_ score is used, and if the blind classifier performs better than the null model with 95 % confidence, then the null model is rejected, and it is concluded that LTM is detected.

#### Behavioural features analysis

In the quantification of STM and LTM by comparing the evolution of the described behavioural features (Table 1) across trials, variation is expressed as the metric’s ratio (*r*). The median statistic is employed to minimize the influence of outliers. Due to differences in the distributions of each metric and the lack of prior information, separate analyses were conducted using non-parametric methods. The metrics evaluated both the presence of STM and LTM as first-order effects, while variations in these memory functions between the Control and DBS groups were assessed as second-order effects. To address potential inflation of the FDR from multiple comparisons, the Benjamini-Hochberg correction was applied. The FDR was controlled at 0.05 for first-order effects and 0.1 for second-order effects, considering the limited sample size and the greater complexity of the second-order analyses.

To compute confidence intervals for STM and LTM evaluation (first-order effect), comparisons are made using paired observations (e.g. a rat in T1 vs the same rat in T4). Exact, distribution-free confidence intervals for *r* are utilised^39^. A paired contrast involves comparing two sets of data, *X* and *Y*, where each set has an equal number of observations, denoted by *n_1_=n_2_=n*, and each element in *X* is uniquely associated with a corresponding element in *Y*. This type of comparison is appropriate for comparing the results of different trials for rats that have undergone the same treatment. For a given metric evaluated in a certain treatment and the control group, with values *x_i_* and *y_i_* respectively, the variable of interest is:

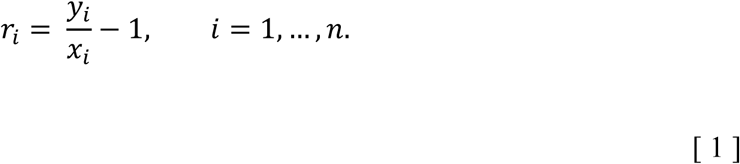

Thus, *r_i_* > 0 (*r_i_* < 0) implies an increase (decrease) in the metric in set Y compared to X with respect to the control group. As observations are paired, the empirical distribution of *r* can be computed and, following Hahn & Meeker^39^, the exact, distribution-free, one-sided confidence lower and upper bound with confidence level 1-*α*, for the quantile *x_p_* are given by the order statistics *x_(l)_* and *x_(u)_*, where:

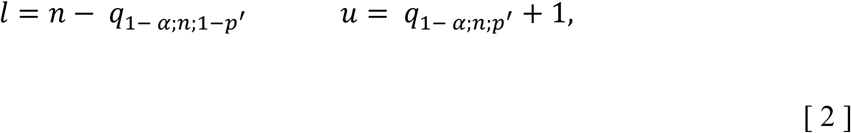

being *q*_*α*;*n*;*p*_the quantile *α* of a binomial distribution *B(n, p)*. The exact, equally-tailed confidence interval with confidence 1-*α* is given by

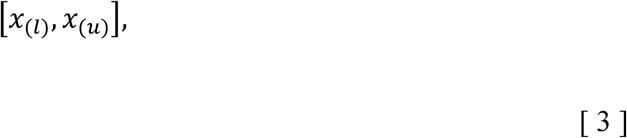

Where *l* and *u* are obtained from [2] using a significance level *α*/2.

When evaluating DBS-ON *vs*. -OFF, the goal is to assess variations in memory (second-order effect). In these cases, pairing individual observations is not feasible. Thus, confidence intervals are computed using a bootstrap method, where pairing occurs at the rat level (block-pairing). Let *X* and *Y* represent the control and treatment groups’ observation sets, which contain subsets associated with each rat:

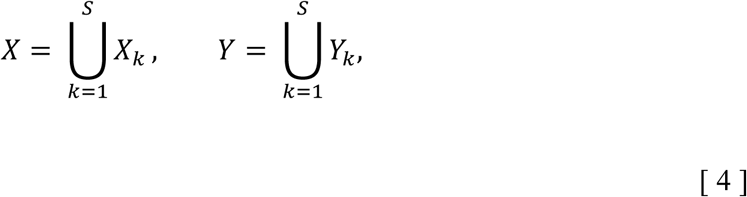

where *S* is the number of rats. Note that the following condition must be met:

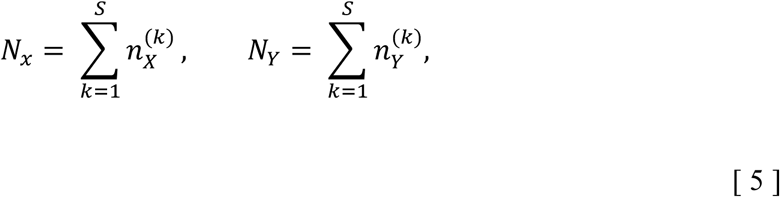

where *N_X_* and *N_Y_* represent the total number of observations for the control and treatment groups, and 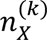 and 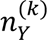 are the number of observations associated with rat *k*.

The procedure involves the following steps:

1. Conduct *R =* 10^4^ resamples with replacement on the paired subsets labelled by the rat (*X_k_, Y_k_*).
2. For each subset *X_k_* and *Y_k_*, resample their contents *R_B_* = 10^3^ times.
3. Calculate the expected values of *X_k_* and *Y_k_*.
4. Compute the variable of interest using the same method as in the paired case with equation [1], using 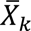 and 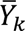 instead of *x_i_* and *y_i_*.
5. Obtain the confidence interval using the standard percentile method with the total computed resamples *R*_*T*_ = *R* · *R*_*B*_ = 10^7^.

Steps 1 and 3 ensure that the conditions for a paired comparison are met, while step 2 accounts for the uncertainty in the expected values.

The original 20 metrics were designed to evaluate the presence of STM and LTM as first-order effects. However, changes in STM and LTM induced by DBS represent second-order effects, building upon the basic presence of memory. From a Bayesian perspective, the likelihood of a metric reliably measuring second-order effects depends on its ability to detect first-order effects. Consequently, to compare memory differences between DBS-ON and DBS-OFF conditions, we included only metrics that were statistically significant in detecting memory presence. Metrics unable to capture the first-order effect, even with a larger sample size, are unlikely to reliably measure second-order effects and were excluded from further analysis. This approach focuses on validated metrics with greater statistical power, reducing the number of tests required and minimizing FDR inflation from multiple comparisons.

#### Conditioned place preference

To investigate the potential appetitive or aversive motivational effects of the stimulation, a CPP protocol was employed. In this paradigm, conditioned preference is assessed by measuring the time spent in the chamber paired with the stimulus before and after conditioning. The protocol spanned three days as follows:

On day 1 animals were allowed to freely explore the apparatus for 15 minutes without any stimulation. The second day the animal was confined to the chamber where it had spent less time in the previous session (non-preferred chamber), and DBS was applied for 30 minutes. On the last day, the animals were once again given the opportunity to freely explore the apparatus without stimulation for 15 minutes.

The proportion of time spent in the conditioned chamber was then quantified to assess the place preference.

#### Elevated plus maze

The EPM serves as a test to assess anxiety levels in experimental animals. The model is based on the natural aversion of rodents to open spaces and their preference for staying in enclosed ones. The test employs an elevated, plus-shaped apparatus with two open arms and two enclosed arms. Anxiety reduction is indicated by an increase in the proportion of time spent in the open arms (time in open arms/total time in open and closed arms).

The experiment comprised a 10-minute duration trial, split into two periods of DBS-ON and two periods of DBS-OFF, each lasting 2 minutes and 30 seconds. These periods were intercalated, randomly assigned, and counterbalanced. The comparison of the time spent by the animals in the open arms versus the closed arms between ON and OFF periods served as an index for evaluating the potential anxiolytic effect of NAc-DBS.

#### Chronic implantation surgery

Following the completion of the behavioural training, the animals underwent bilateral implantation of bipolar custom-made electrodes in the NAc. The rats were anaesthetised using isoflurane (induction at 5 %, maintenance at 1.5-2.5 % in oxygen) and received local analgesia through a lidocaine injection (50 mg/ml). Positioned in a stereotaxic frame (Narishige, Japan), the animals’ temperature was maintained at 37 °C using a heat blanket (Cibertec, Spain), and heart and breath rates were monitored to adjust the isoflurane concentration (MoseOx Plus, Starr Life Sciences, USA). Ophthalmic gel was applied to keep the animal’s eyes hydrated throughout the procedure.

Electrodes were constructed by twisting two teflon-coated platinum-iridium filaments (A-M Systems, USA), resulting in 200 μm in diameter electrodes, with an impedance of 80-100 kΩ. Two of these electrodes were soldered to a Lemo connector (Fig. S1A, 1S series, Lemo, Switzerland) to connect to the pulse generator and current source (STG-2004, Multi-Channel Systems, Germany) through a custom-made cable (Fig. S1A). Electrodes were targeted at both NAc through trephines drilled in the skull. The target coordinates were anteroposterior (AP) 1.5 mm from bregma, mediolateral (ML) 0.8 mm, and dorsoventral (DV) 6.5 mm from dura (Fig. 3H), following the Paxinos and Watson atlas^40^. To ensure implant durability, two screws were affixed to the skull, and the implant secured in place with dental cement (Superbond, Sun Medical, Japan).

Towards the end of the procedure, animals received a buprenorphine injection (3 μg/kg). For the subsequent three days, or until signs of discomfort ceased, they were provided with pellets infused with buprenorphine (0.003 mg/ml). After three days, the animals were rehoused in pairs and allowed to recover for a minimum of 7 days before any behavioural testing continued.

#### MRI-compatible surgery

Functional MRI experiments were performed under urethane anaesthesia (1.3 g/kg, i.p.), with oxygen supplied during surgical procedures as previously described. To minimise artefacts during electrical stimulation inside the scanner, glass-coated carbon-fibre bipolar electrodes developed in the laboratory were employed as previously described^41,42^. Individual carbon fibres with a diameter of 7 μm (Goodfellow Cambridge Limited, UK) were inserted into bundles within a theta-shaped glass capillary (World Precision Instruments) previously pulled from 7 mm long pipettes. This configuration resulted in a tip of 200 μm and was adjusted to achieve an electrical impedance of 40-65 kΩ. A regular wire with a gold pin connector was attached to the pipette, connected to the carbon fibres using silver conductive epoxy resin (RS Components, UK), and isolated with clear epoxy resin. Subsequently, the tip was bent in a flame to a 90° angle to minimise implant height, allowing for close proximity of the MRI array coil and the animal’s brain (Fig. S1B). This preparation has been shown to provide robust fMRI results in rats^43^.

These carbon-fibre electrodes were inserted into the NAc at coordinates 1.2 mm AP, 0.8 mm ML, and 6.6 mm DV (Fig. 4A). The electrode was firmly secured in place with MRI-compatible dental cement (Heraeus Medical, Wehrheim, Germany), and the animal was then transferred to the scanner.

#### fMRI experiments and data analysis

For the fMRI experiments, the previously prepared urethane-anaesthetised animals were positioned in a custom-made animal holder with adjustable bite and ear bars, and then placed on the magnet bed. A constant supply of 0.8 l/min O2 was provided, and the temperature was maintained between 37 and 37.5°C using a water heat pad. Temperature, heart rate, and SpO2 were monitored as during surgeries. The experiments were conducted in a horizontal 7 T scanner with a 30 cm diameter bore (Biospec 70/30, Bruker Medical, Ettlingen, Germany). A 1H rat brain receive-only phased-array coil with an integrated combiner and preamplifier, and no-tune/no-match, was utilised along with the actively detuned transmit-only resonator (Bruker BioSpin MRI GmbH, Germany).

Acquisition involved 15 coronal slices using a gradient echo-echo planar image (GE-EPI) sequence with the following parameters: field of view (FOV) 25 x 25 mm, slice thickness 1 mm, matrix 96 x 96, segments 1, flip angle (FA) 60°, time echo (TE) 15 ms, time repetition (TR) 2 s. For the acquisition of the 130 Hz trains, a total of 150 volumes were acquired over a 5-minute duration, and 3 repetitions of this protocol were acquired per animal for averaging purposes. In the continuous stimulation paradigm, 4 repetitions of 150 volumes were acquired in the middle of the 30-minute stimulation sequence (see Fig. 4B). Similar acquisitions were conducted before any stimulation to establish a baseline record of activity in the absence of stimulation.

T2-weighted anatomical images were obtained using a rapid acquisition relaxation enhanced sequence (RARE) with parameters: FOV 25 x 25 mm, 15 slices, slice thickness 1 mm, matrix 192 x 192, TEeff 56 ms, TR 2 s, RARE factor 8.

fMRI data were analysed offline using custom software developed in Python, which included the Advanced Normalization Tools Ecosystem (ANTs, https://github.com/ANTsX), FSL Software (https://fsl.fmrib.ox.ac.uk/fsl/fslwiki/FSL), and Analysis of Functional Neuroimage (AFNI, https://afni.nimh.nih.gov/).

The images were augmented 10 times, and anatomical ones were utilised to create a common template for brain extraction mask generation. Anatomical images underwent bias field correction and were registered to the custom template for brain extraction. The preprocessing of functional images included motion correction (McFlirt), brain extraction with registration to the template, intensity normalisation, and temporal (high-pass 0.01 Hz) and spatial filtering (FWHM with a kernel ∼1.5x voxel size).

Activation maps were processed using FEAT (FSL’s FMRI Expert Analysis Tool). A time series representing each of the stimulation trains was convolved with a double-gamma hemodynamic response function shifted forward in time by 2 seconds, which was then used for the general linear model analysis. Time-series statistical analysis was conducted using FILM (FMRIB’s Improved Linear Model) with local autocorrelation correction^44^. Higher-level analysis was performed using a fixed-effects model for averaging between runs, and a mixed-effects model utilising FLAME (FMRIB’s Local Analysis of Mixed Effects) with automatic outlier detection^45-47^ for subjects averaging. z (Gaussianised T/F) statistical images were thresholded using clusters determined by z > 2.33 and a corrected cluster significance of p = 0.05^48^.

The BOLD time series from regions of interest were extracted by registering the subjects to an anatomical atlas^49^. This registration facilitated the calculation of the mean BOLD response in those regions triggered by each stimulation train as a percentage relative to a peristimulus baseline of 8s. Additionally, the Pearson correlation between different structures was computed using the BOLD time series from the registered ROIs in both resting and stimulation conditions.

#### Histology

To verify the accurate placement of implanted elements and assess potential excessive tissue damage, immunohistochemical analyses were conducted.

Following the completion of experiments, the animals underwent transcardial perfusion with a phosphate saline buffer at 37 °C, followed by 4% paraformaldehyde at 4 °C. Subsequently, the fixed brains were extracted from the skulls and immersed in paraformaldehyde at room temperature overnight. Coronal slices of 100 μm thickness were obtained using a vibratome (Leica Biosystems, Germany).

For labelling glial fibrillary acidic protein, a mouse monoclonal antibody (Sigma Aldrich, USA) was utilised, followed by a secondary anti-mouse Alexa Fluor 488-conjugated antibody (Thermo Fisher Scientific, USA). Slices were further stained with 4’,6-diamino-2-phenylidole. Fluorescence microscope images were captured (Leica Microsystems, Germany) to visualise the location of the cannulas and the glial reaction. Image analysis was performed using the ImageJ software^50^.

---

**Fig. S1.**
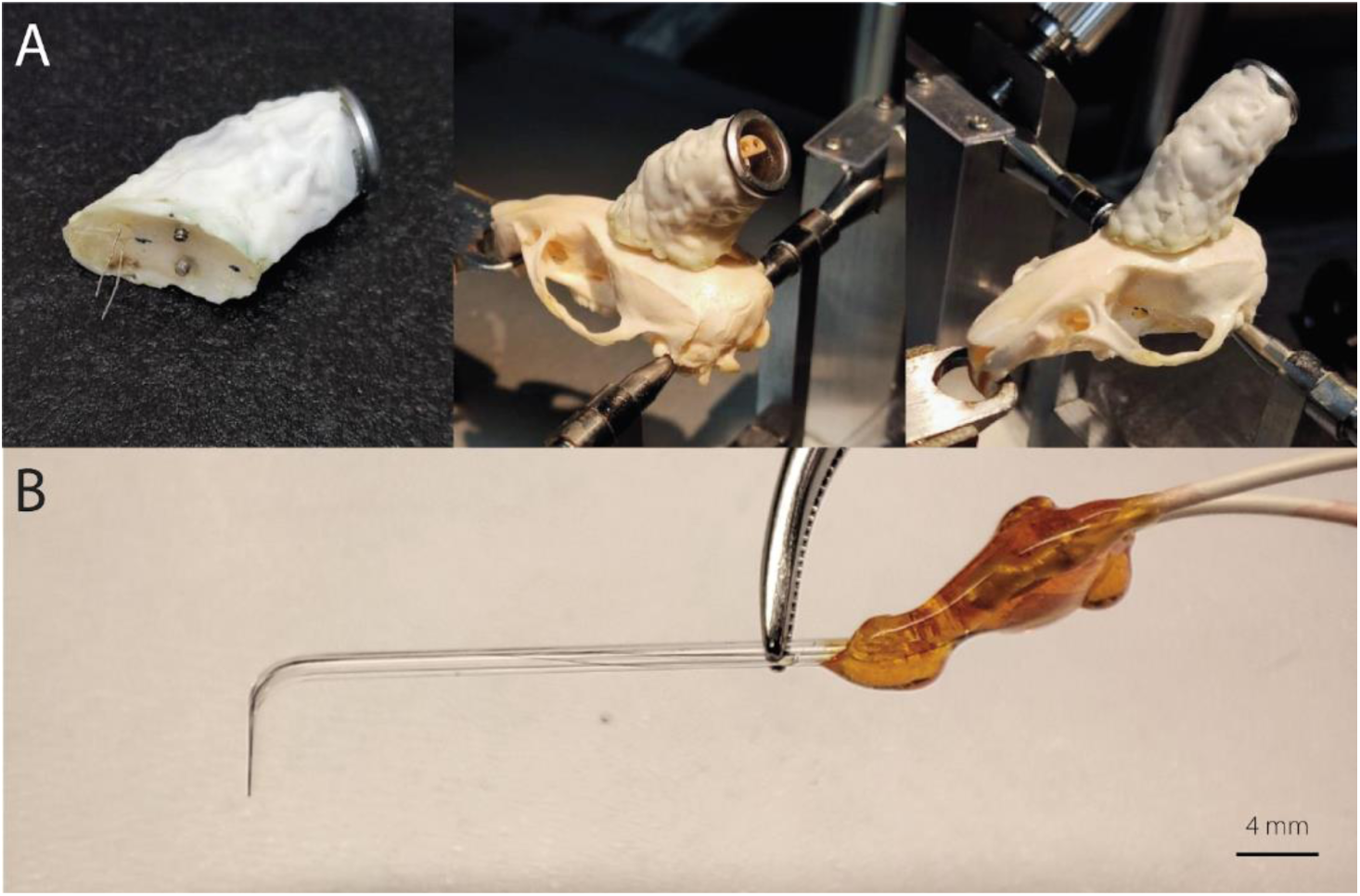
DBS implants. **(A)** Behavioural chronic implant with bipolar bilateral Pt-Ir electrodes and its location over a rat skull. **(B)** Borosilicate-carbon fibre bipolar MRI-compatible electrode.

**Fig. S2.**
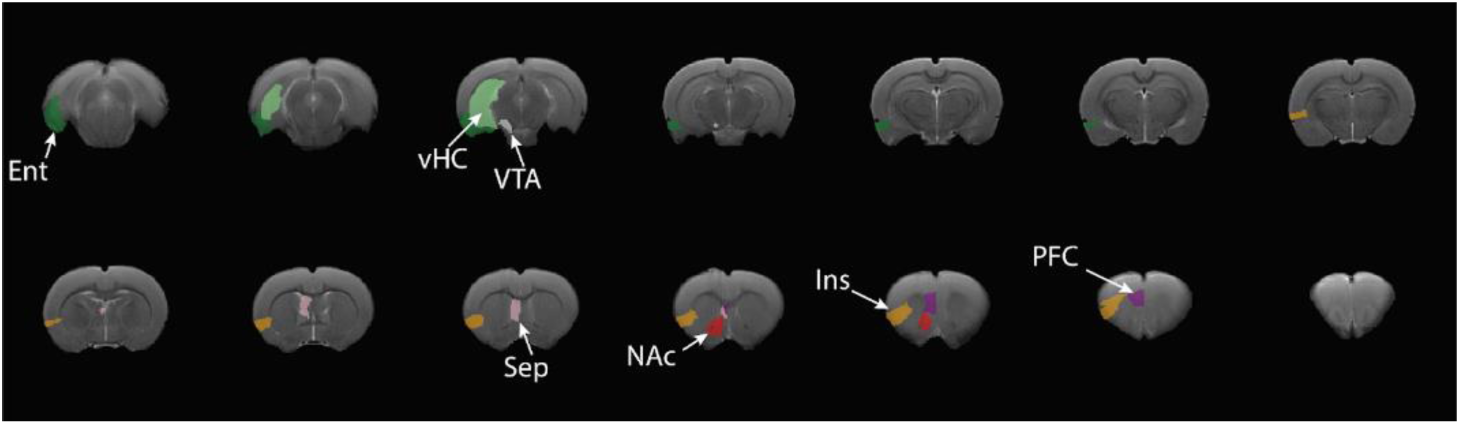
ROIs for mean BOLD extraction. NAc: nucleus accumbens (red), PFC: prefrontal cortex (purple), Ins: insular cortex (orange), Ent: entorhinal cortex (dark green), Sep: septum (pink), vHC: ventral hippocampus (light green), VTA: ventral tegmental area (white).

**Table S1.**
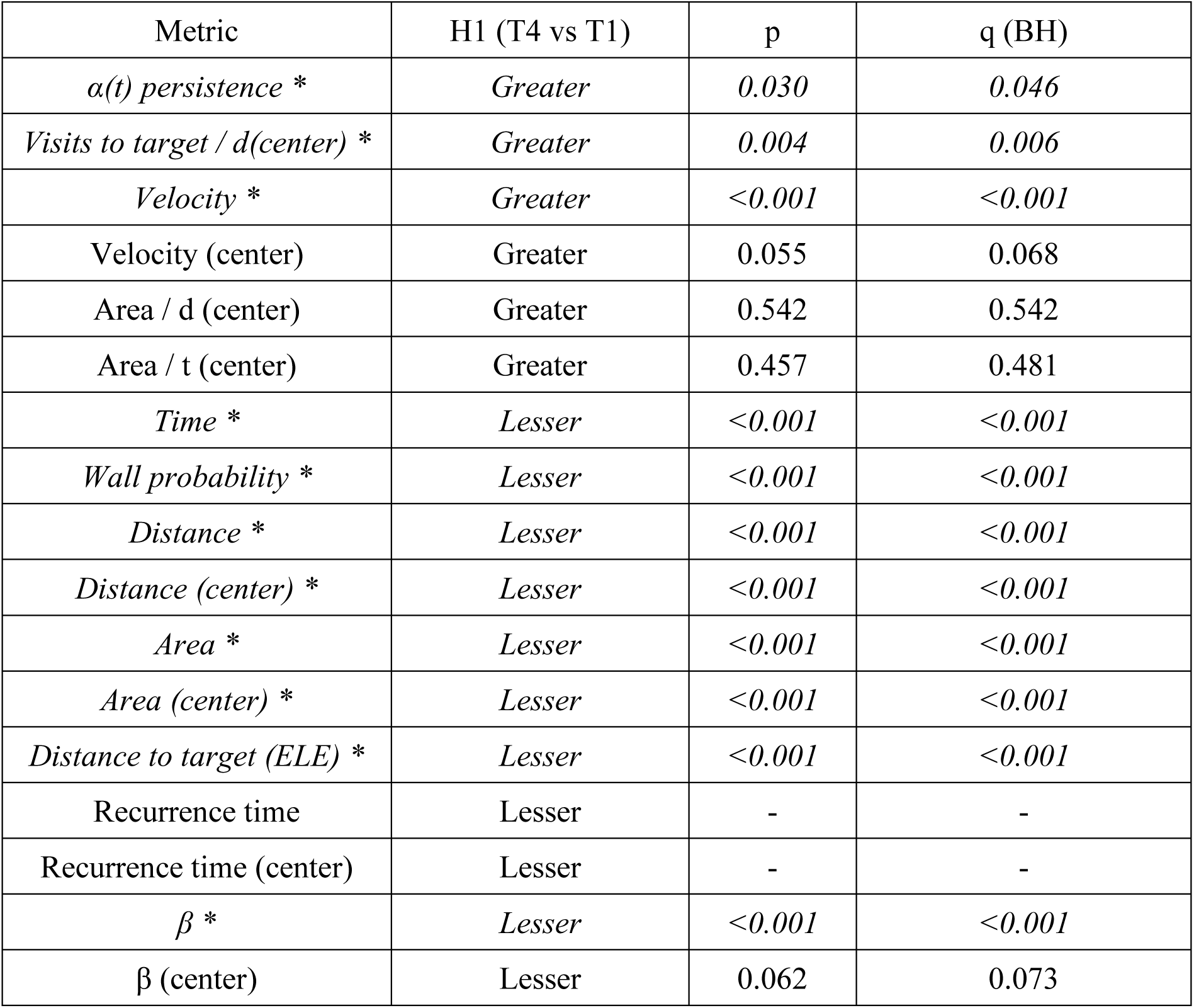
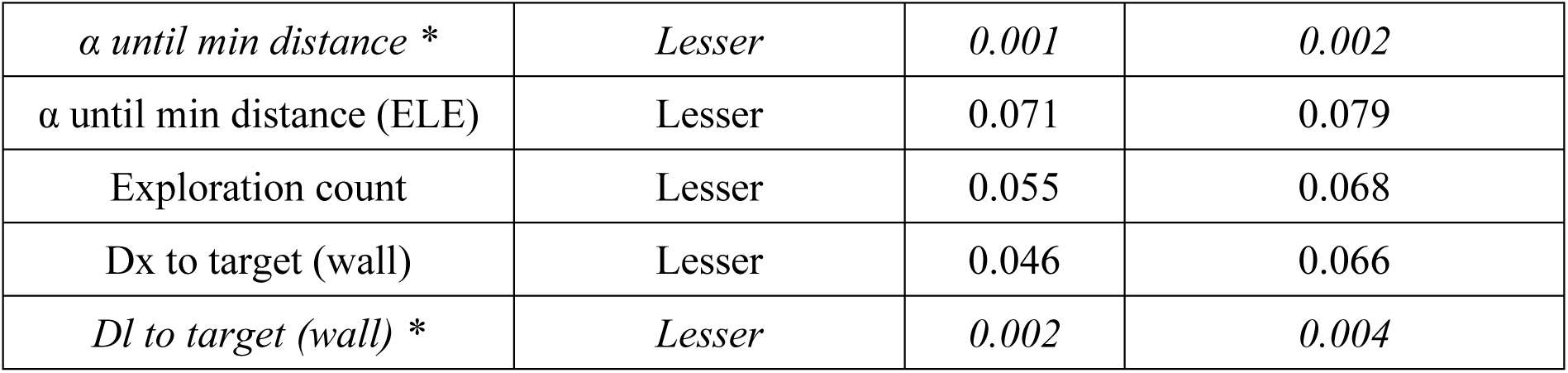
Selection of STM metrics based on effect detection in the memory evaluation group. The q-values represent p-values adjusted for multiple comparisons, controlling the false discovery rate (FDR) using the Benjamini-Hochberg method. Metrics with q < FDR = 0.05 are in italic and marked with an asterisk (*). A hyphen indicates insufficient data (n < 2).

**Table S2.**
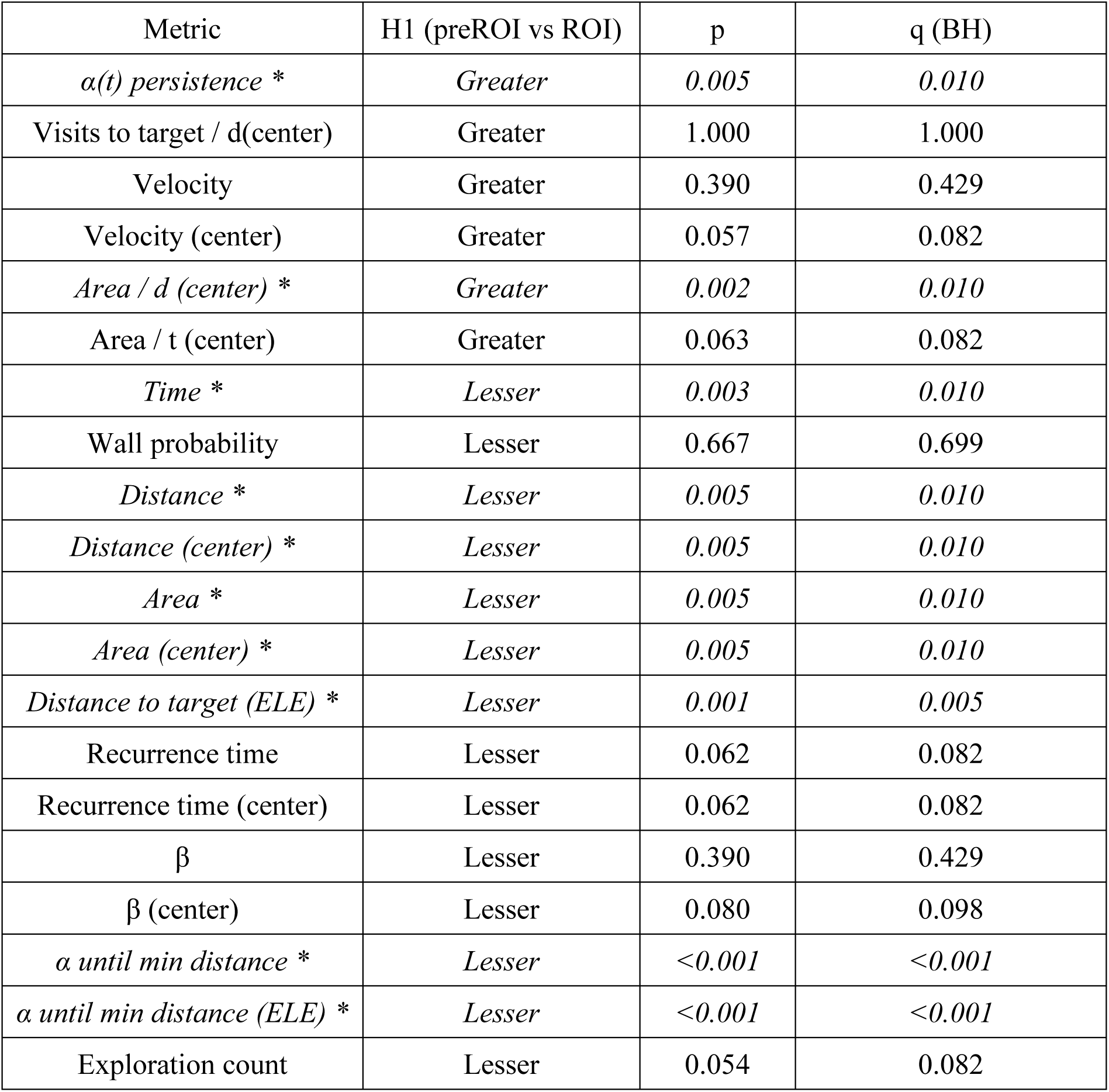
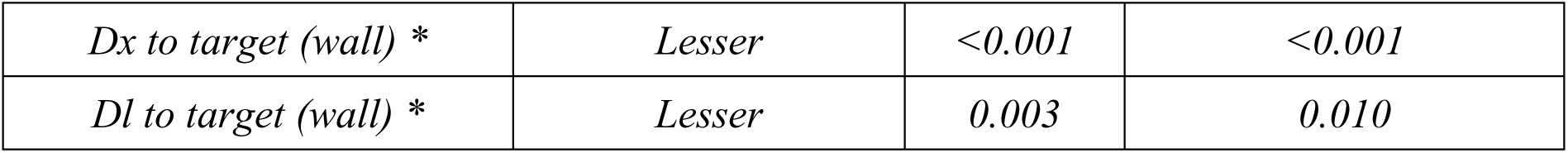
Selection of LTM metrics based on effect detection in the first-order spatial memory analysis. The q-values represent p-values adjusted for multiple comparisons, controlling the false discovery rate (FDR) using the Benjamini-Hochberg method. Metrics with q < FDR = 0.05 are in italic and marked with an asterisk (*).

**Table S3.**
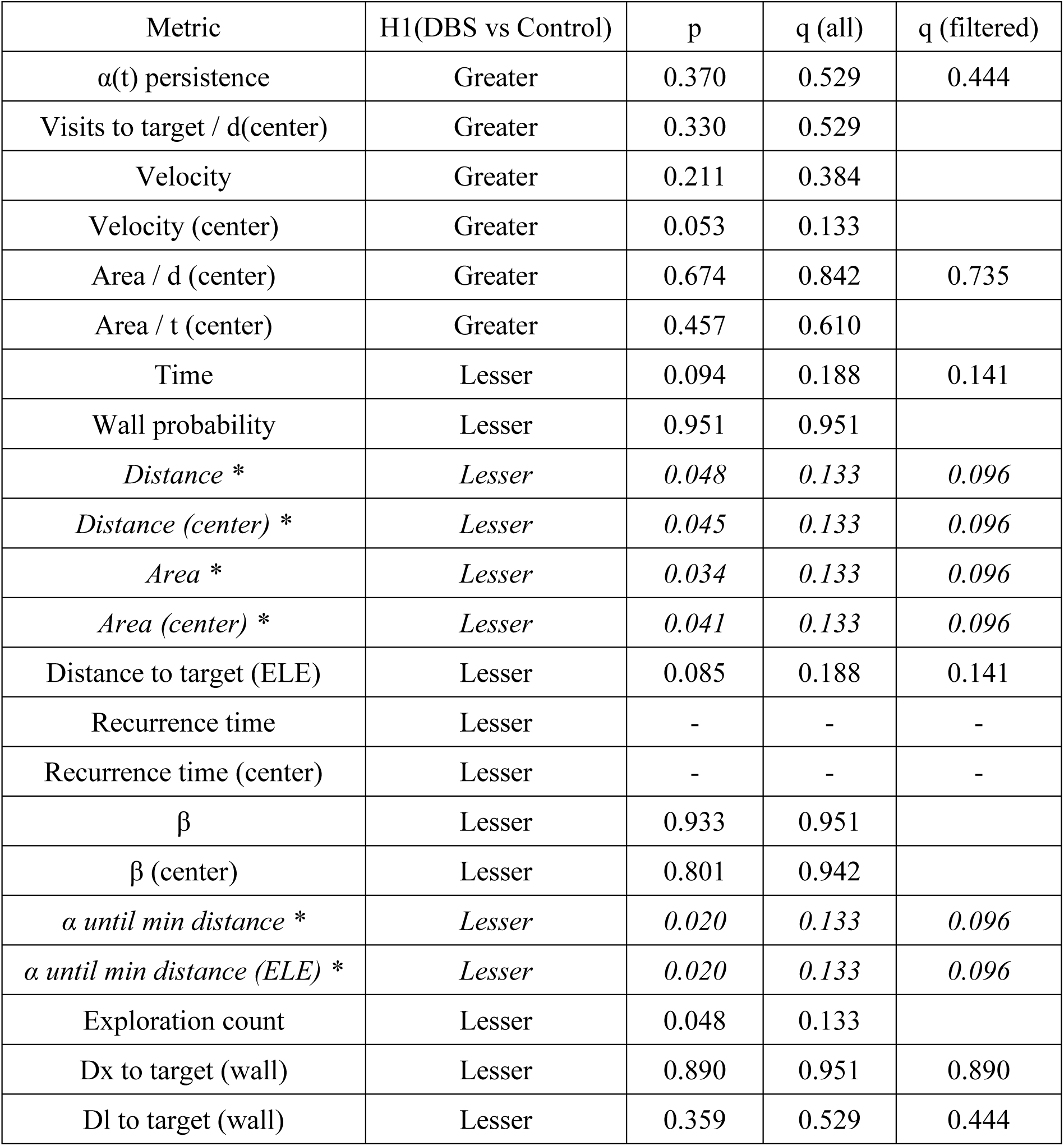
Effects of DBS on LTM encoding. The q-values represent p-values adjusted for multiple comparisons, controlling the false discovery rate (FDR) using the Benjamini-Hochberg method. q (all) refers to calculations considering all metrics, while q (filtered) includes only metrics that detected LTM in the first-order spatial memory analysis (Table S2). Metrics with q (filtered) < FDR = 0.10 are in italic and marked with an asterisk (*). A hyphen indicates insufficient data (n < 2).

## AVAILABILITY OF DATA AND MATERIALS

All data will be available at DIGITAL.CSIC repository.

## Notes

### Competing Interest Statement

The authors have declared no competing interest.

