## Supplementary Information for "NAc-DBS selectively enhances memory updating without effect on retrieval"

### Supplementary methods

#### *Tracking analysis*

DeepLabCut software[21,22] was used to extract the precise position of the selected body parts of the animals across each frame of the videos. Each condition (wired and non-wired animals) involved preparing two sets of 16 videos for training. For each set, 40 frames of each video were utilised to label desired body parts (nose, left and right ears, shoulders, back, and the base of the tail, Fig.1B). Subsequently, 95% of the labelled frames were used for training the network.

A ResNet-101-based neural network[37,38] was employed with default parameters for 1.3 x $10^6$ training iterations and three consecutive training sessions. The results were validated through a single shuffle, yielding a test error of 3.95 pixels and a train error of 3.5 pixels for the control condition. In the DBS condition, the test error was 5.17 pixels, and the train error was 3.9 pixels. The image size used was 1552 x 1552 pixels. Subsequently, these networks were applied to analyse the remaining recorded videos for each respective group.

#### *Long-term memory evaluation*

For LTM detection and regarding the probability of a rat in finding the target ROI or any other ROI in the arena, we defined a null model (lack of LTM): a binary classifier predicting that a rat first finds the target (either ROI or previous day's ROI) whose location is closer to the rat's entering point to the arena:

$$X = (x_1, \ldots, x_n), \qquad x_i = \begin{cases} 1, & d(rat, p.ROI) < d(rat, ROI), \\ 0, & d(rat, p.ROI) > d(rat, ROI), \end{cases}$$

The null model is then evaluated against the actual outcome (finding first the previous day's ROI or the ROI at each T1) across rats and days:

$$Y = (y_1, \dots, y_n), \qquad y_i = \begin{cases} 1, & if\ rat\ finds\ preROI, \\ 0, & if\ rat\ finds\ ROI. \end{cases}$$

Lack of LTM suggests that this null model should outperform a blind classifier that simply predicts the majority class (without using any data). To compare both classifiers, the $F_1$ score is used, and if the blind classifier performs better than the null model with 95 % confidence, then the null model is rejected, and it is concluded that LTM is detected.

$$l = n - q_{1-\alpha;n;1-p'} \qquad u = q_{1-\alpha;n;p'} + 1,$$

[ 2 ]

being $q_{\alpha;n;p}$ the quantile $\alpha$ of a binomial distribution $B(n, p)$. The exact, equally-tailed confidence interval with confidence 1-$\alpha$ is given by

$$\left[x_{(l)}, x_{(u)}\right],$$

[ 3 ]

Where $l$ and $u$ are obtained from [2] using a significance level $\alpha/2$.

When evaluating DBS-ON *vs*. -OFF, the goal is to assess variations in memory (second-order effect). In these cases, pairing individual observations is not feasible. Thus, confidence intervals are computed using a bootstrap method, where pairing occurs at the rat level (block-pairing). Let $X$ and $Y$ represent the control and treatment groups' observation sets, which contain subsets associated with each rat:

$$X = \bigcup_{k=1}^{S} X_k \,, \qquad Y = \bigcup_{k=1}^{S} Y_k,$$

[ 4 ]

where $S$ is the number of rats. Note that the following condition must be met:

$$N_x = \sum_{k=1}^{S} n_X^{(k)} \,, \qquad N_Y = \sum_{k=1}^{S} n_Y^{(k)},$$

[ 5 ]

where $N_X$ and $N_Y$ represent the total number of observations for the control and treatment groups, and $n_X^{(k)}$ and $n_Y^{(k)}$ are the number of observations associated with rat $k$.

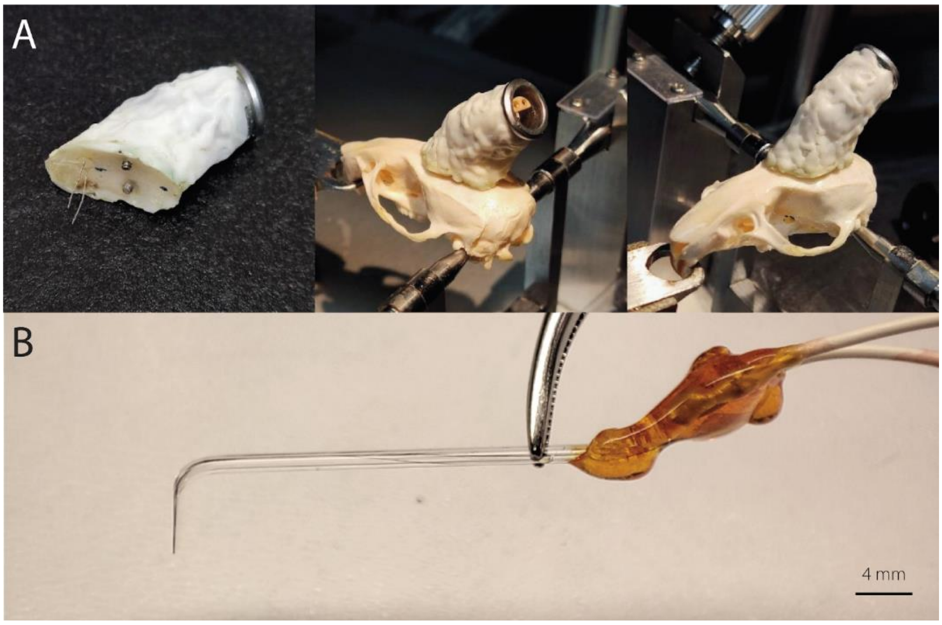

**Fig. S1. DBS implants. (A)** Behavioural chronic implant with bipolar bilateral Pt-Ir electrodes and its location over a rat skull. **(B)** Borosilicate-carbon fibre bipolar MRI-compatible electrode.

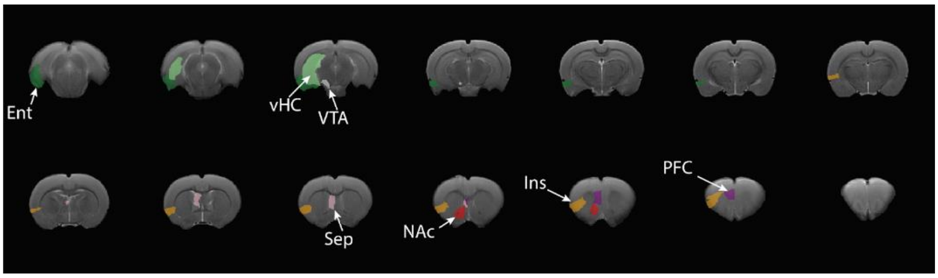


**Fig. S2. ROIs for mean BOLD extraction.** NAc: nucleus accumbens (red), PFC: prefrontal cortex (purple), Ins: insular cortex (orange), Ent: entorhinal cortex (dark green), Sep: septum (pink), vHC: ventral hippocampus (light green), VTA: ventral tegmental area (white).

| Metric | H1 (T4 vs T1) | p | q (BH) |
|---|---|---|---|
| α(t) persistence * | Greater | 0.030 | 0.046 |
| Visits to target / d(center) * | Greater | 0.004 | 0.006 |
| Velocity * | Greater | <0.001 | <0.001 |
| Velocity (center) | Greater | 0.055 | 0.068 |
| Area / d (center) | Greater | 0.542 | 0.542 |
| Area / t (center) | Greater | 0.457 | 0.481 |
| Time * | Lesser | <0.001 | <0.001 |
| Wall probability * | Lesser | <0.001 | <0.001 |
| Distance * | Lesser | <0.001 | <0.001 |
| Distance (center) * | Lesser | <0.001 | <0.001 |
| Area * | Lesser | <0.001 | <0.001 |
| Area (center) * | Lesser | <0.001 | <0.001 |
| Distance to target (ELE) * | Lesser | <0.001 | <0.001 |
| Recurrence time | Lesser | - | - |
| Recurrence time (center) | Lesser | - | - |
| β * | Lesser | <0.001 | <0.001 |
| β (center) | Lesser | 0.062 | 0.073 |

| | | | |
|---|---|---|---|
| *α until min distance* * | *Lesser* | *0.001* | *0.002* |
| α until min distance (ELE) | Lesser | 0.071 | 0.079 |
| Exploration count | Lesser | 0.055 | 0.068 |
| Dx to target (wall) | Lesser | 0.046 | 0.066 |
| *Dl to target (wall)* * | *Lesser* | *0.002* | *0.004* |

**Table S1. Selection of STM metrics based on effect detection in the memory evaluation group.** The q-values represent p-values adjusted for multiple comparisons, controlling the false discovery rate (FDR) using the Benjamini-Hochberg method. Metrics with q < FDR = 0.05 are in italic and marked with an asterisk (*). A hyphen indicates insufficient data (n < 2).

| Metric | H1 (preROI vs ROI) | p | q (BH) |
|---|---|---|---|
| *α(t) persistence* * | *Greater* | *0.005* | *0.010* |
| Visits to target / d(center) | Greater | 1.000 | 1.000 |
| Velocity | Greater | 0.390 | 0.429 |
| Velocity (center) | Greater | 0.057 | 0.082 |
| *Area / d (center)* * | *Greater* | *0.002* | *0.010* |
| Area / t (center) | Greater | 0.063 | 0.082 |
| *Time* * | *Lesser* | *0.003* | *0.010* |
| Wall probability | Lesser | 0.667 | 0.699 |
| *Distance* * | *Lesser* | *0.005* | *0.010* |
| *Distance (center)* * | *Lesser* | *0.005* | *0.010* |
| *Area* * | *Lesser* | *0.005* | *0.010* |
| *Area (center)* * | *Lesser* | *0.005* | *0.010* |
| *Distance to target (ELE)* * | *Lesser* | *0.001* | *0.005* |
| Recurrence time | Lesser | 0.062 | 0.082 |
| Recurrence time (center) | Lesser | 0.062 | 0.082 |
| β | Lesser | 0.390 | 0.429 |
| β (center) | Lesser | 0.080 | 0.098 |
| *α until min distance* * | *Lesser* | *<0.001* | *<0.001* |
| *α until min distance (ELE)* * | *Lesser* | *<0.001* | *<0.001* |
| Exploration count | Lesser | 0.054 | 0.082 |

| | | | |
|---|---|---|---|
| *Dx to target (wall) ** | *Lesser* | *<0.001* | *<0.001* |
| *Dl to target (wall) ** | *Lesser* | *0.003* | *0.010* |

**Table S2. Selection of LTM metrics based on effect detection in the first-order spatial memory analysis.** The q-values represent p-values adjusted for multiple comparisons, controlling the false discovery rate (FDR) using the Benjamini-Hochberg method. Metrics with q < FDR = 0.05 are in italic and marked with an asterisk (*).

| Metric | H1(DBS vs Control) | p | q (all) | q (filtered) |
|---|---|---|---|---|
| α(t) persistence | Greater | 0.370 | 0.529 | 0.444 |
| Visits to target / d(center) | Greater | 0.330 | 0.529 | |
| Velocity | Greater | 0.211 | 0.384 | |
| Velocity (center) | Greater | 0.053 | 0.133 | |
| Area / d (center) | Greater | 0.674 | 0.842 | 0.735 |
| Area / t (center) | Greater | 0.457 | 0.610 | |
| Time | Lesser | 0.094 | 0.188 | 0.141 |
| Wall probability | Lesser | 0.951 | 0.951 | |
| *Distance ** | *Lesser* | *0.048* | *0.133* | *0.096* |
| *Distance (center) ** | *Lesser* | *0.045* | *0.133* | *0.096* |
| *Area ** | *Lesser* | *0.034* | *0.133* | *0.096* |
| *Area (center) ** | *Lesser* | *0.041* | *0.133* | *0.096* |
| Distance to target (ELE) | Lesser | 0.085 | 0.188 | 0.141 |
| Recurrence time | Lesser | - | - | - |
| Recurrence time (center) | Lesser | - | - | - |
| β | Lesser | 0.933 | 0.951 | |
| β (center) | Lesser | 0.801 | 0.942 | |
| *α until min distance ** | *Lesser* | *0.020* | *0.133* | *0.096* |
| *α until min distance (ELE) ** | *Lesser* | *0.020* | *0.133* | *0.096* |
| Exploration count | Lesser | 0.048 | 0.133 | |
| Dx to target (wall) | Lesser | 0.890 | 0.951 | 0.890 |
| Dl to target (wall) | Lesser | 0.359 | 0.529 | 0.444 |

**Table S3. Effects of DBS on LTM encoding.** The q-values represent p-values adjusted for multiple comparisons, controlling the false discovery rate (FDR) using the Benjamini-Hochberg method. q (all) refers to calculations considering all metrics, while q (filtered) includes only metrics that detected LTM in the first-order spatial memory analysis (Table S2). Metrics with q (filtered) < FDR = 0.10 are in italic and marked with an asterisk (*). A hyphen indicates insufficient data (n < 2).
